# Multi-genomic analysis of the Cation Diffusion Facilitator transporters from algae

**DOI:** 10.1101/2019.12.18.880724

**Authors:** Aniefon Ibuot, Andrew P. Dean, Jon K. Pittman

**Author notes:** Corresponding Author: Dr Jon Pittman, Department of Earth and Environmental Sciences, The University of Manchester, Michael Smith Building, Oxford Road, Manchester M13 9PT, UK.

## Abstract

Metal transport processes are relatively poorly understood in algae in comparison to higher plants and other eukaryotes. A screen of genomes from 33 taxonomically diverse algal species was conducted to identify members of the Cation Diffusion Facilitator (CDF) family of metal ion transporter. All algal genomes contained at least one CDF gene with four species having >10 CDF genes (median of 5 genes per genome), further confirming that this is a ubiquitous gene family. Phylogenetic analysis suggested a CDF gene organisation of five groups, which includes Zn-CDF, Fe/Zn-CDF and Mn-CDF groups, consistent with previous phylogenetic analyses, and two functionally undefined groups. One of these undefined groups was algal specific although excluded chlorophyte and rhodophyte sequences. The majority of sequences (22 out of 26 sequences) from this group had a putative ion binding site motif within transmembrane domain 2 and 5 that was distinct from other CDF proteins, such that alanine or serine replaced the conserved histidine residue. The phylogenetic grouping was supported by sequence cluster analysis. Yeast heterologous expression of CDF proteins from *Chlamydomonas reinhardtii* indicated Zn^2+^ and Co^2+^ transport function by CrMTP1, and Mn^2+^ transport function by CrMTP2, CrMTP3 and CrMTP4, which validated the phylogenetic prediction. However, the Mn-CDF protein CrMTP3 was also able to provide zinc and cobalt tolerance to the Zn- and Co-sensitive *zrc1cot1* yeast strain. There is wide diversity of CDF transporters within the algae lineage, and some of these genes may be attractive targets for future applications of metal content engineering in plants or microorganisms.

## Introduction

Knowledge of the transport pathways for metal influx, efflux and subcellular metal partitioning in plant and algal cells is critical in order to understand the processes of essential metal homeostasis within the cell. These will include pathways for essential trace metals such as zinc, copper and manganese, which are required for many biochemical functions including as enzyme co-factors or structural components of proteins (Andreini et al. 2008). Furthermore, both essential and non-essential metals, such as cadmium, are toxic at elevated concentrations and so transporters play a key role in protecting the cell against metal toxicity (Kramer et al. 2007). This protection can be mediated by metal efflux from the cell or intracellular metal sequestration into compartments such as the vacuole (Sharma et al. 2016). Manipulation of expression of specific transporter proteins, such as by genetic engineering, can therefore be used to enhance metal tolerance of an organism (Antosiewicz et al. 2014). Moreover, manipulation of metal sequestration may also provide increased total metal content within a cell whilst still providing metal tolerance, and would be useful for metal bioremediation applications (Cheng et al. 2019; Ibuot et al. 2017).

Algae are one such group of organisms that have been proposed as potentially suitable for metal bioremediation (Suresh Kumar et al. 2015), but if genetic engineering of metal transporters is to be tested in algae then molecular targets that have been functionally characterised are needed. Furthermore, knowledge of the gene families that encode metal transporters across taxonomically diverse algal species provides understanding of the evolution of metal homeostasis within these important photosynthetic organisms. There is a long history of the study of metal homeostasis and transport processes in algae during the second half of the last century (particularly for green microalgae species and brown seaweeds), including physiological investigation of metal micronutrient requirements and metal toxicity responses, and biochemical characterisation of metal transport (Fig. S1). However, the molecular identification of individual metal transporters and genomic scale analysis of whole gene families has only begun in the last fifteen years since the availability of algal genome sequences. For example, genomic analyses of green microalgae, such as *Chlamydomonas reinhardtii*, and red microalgae, such as *Cyanidioschyzon merolae* indicate that there are a wide variety of metal transporter pathways that are conserved between microalgae and higher plants alongside other eukaryotes such as yeast (Hanikenne et al. 2005; Blaby-Haas and Merchant 2012). The identification of putative algal metal transporter genes has been complemented by other genome-scale analyses such as by proteomic and transcriptomic methods (Malasarn et al. 2013; Beauvais-Flück et al. 2019; Puente-Sánchez et al. 2018; Hsieh et al. 2013; Khatiwada et al. 2020), and subsequent functional characterisation of individual metal transporters (Ibuot et al. 2017; Tsednee et al. 2019; Chang et al. 2020).

One important metal transporter family is the Cation Diffusion Facilitator (CDF) family, which are referred to as Metal Transport Proteins (MTP) in higher plants (Kolaj-Robin et al. 2015; Ricachenevsky et al. 2013). CDF-type transporters have been reported to facilitate the transport of metals including cadmium (Cd), cobalt (Co), copper (Cu), iron (Fe), manganese (Mn), nickel (Ni), and zinc (Zn), and are found virtually in all organisms covering the Eukaryote, Eubacteria and Archaea kingdoms (Cubillas et al. 2013; Nies and Silver 1995). In yeast and plants, CDF/MTP proteins localise to membranes of various intracellular compartments, in particular the vacuolar membrane but also the mitochondria, endoplasmic reticulum (ER), Golgi/trans-Golgi and pre-vacuolar compartment, where they are involved in the efflux of metals from the cytoplasm into these organelles (Haney et al. 2005; Ricachenevsky et al. 2013). For example, in *Saccharomyces cerevisiae*, CDF proteins mediate Zn^2+^ and Co^2+^ transfer into the vacuole (ScZrc1 and ScCot1), Zn^2+^ transfer into the ER (ScZrg17 and ScMsc2) and Fe^2+^ transfer from the mitochondria (ScMmt1 and ScMmt2) (Ellis et al. 2005; Li et al. 2014; MacDiarmid et al. 2002). In *Arabidopsis thaliana*, AtMTP1 and related orthologues including AtMTP3 play a major role in the storage and sequestration of Zn^2+^ in the vacuole (Arrivault et al. 2006; Kobae et al. 2004), AtMTP5 and AtMTP12 transport Zn^2+^ into the Golgi (Fujiwara et al. 2015), while AtMTP8 and AtMTP11 mediate Mn^2+^ transport into the vacuole and Golgi (Delhaize et al. 2007; Peiter et al. 2007; Eroglu et al. 2016).

These main metal substrates (Zn^2+^, Co^2+^, Mn^2+^, Fe^2+^) of the higher plant and fungal CDF family proteins are all essential micronutrients but can cause major toxicity if they accumulate at excess concentration. Zn is an important co-factor for many enzymes, such as carbonic anhydrase that is required for photosynthetic carbon fixation, and is associated with many Zn-finger containing transcription factors, and has a structural role with some proteins (Decaria et al. 2010; Hänsch and Mendel 2009). Co is an essential metal in a few organisms, such as where it is required for vitamin B_12_ and used by coenzyme B_12_-dependent enzymes (Renfrew et al. 2018). Mn also has many essential enzyme co-factor requirements, such as Mn-dependent superoxide dismutase and the water splitting oxygen evolution process of Photosystem II (Pittman 2005; Socha and Guerinot 2014). Finally, Fe also plays many co-factor roles and protein structural roles, and is essential for electron transport processes (Hänsch and Mendel 2009).

The current understanding of CDF/MTP proteins in algae is limited. The CDF gene family appears to be present in all previously examined algal species but specific details about CDF family structures within individual species are unknown (Blaby-Haas and Merchant 2012, 2017; Gustin et al. 2011). Phylogenetic analysis will be able to reveal some useful details since substrate specificity seems to be related to phylogeny (Cubillas et al. 2013; Montanini et al. 2007). Within the well-studied model microalga *C. reinhardtii* there are five putative MTP genes (CrMTP1 to CrMTP5). Some of these genes are transcriptionally induced by Zn or Mn deficiency conditions (Allen et al. 2007; Malasarn et al. 2013), but to date only CrMTP4 has been functionally characterised as a Mn^2+^ and Cd^2+^ transporter (Ibuot et al. 2017). This was determined by performing over-expression of CrMTP4 in *C. reinhardtii* yielding enhanced Cd tolerance, and by using yeast (*S. cerevisiae*) heterologous expression where CrMTP4 was able to rescue the Mn sensitive phenotype of the *pmr1* yeast mutant. In this study, we aimed to perform a large-scale phylogenetic analysis of putative MTP genes from available algal genome sequences in order to evaluate the potential diversity within this broad taxonomic lineage. Furthermore, a characterisation of additional members of the *C. reinhardtii* MTP gene family was carried out by using yeast heterologous expression to infer the metal substrates and determine metal tolerance functions.

## Materials and Methods

### Bioinformatic analysis

Thirty three algal species genomes were screened from one Charophyta: *Klebsormidium nitens*; 13 Chlorophyta: *Bathycoccus prasinos* RCC1105, *Botryococcus braunii*, *Chlamydomonas reinhardtii*, *Chlorella* sp. NC64A, *Chromochloris zofingiensis*, *Coccomyxa subellipsoidea* C-169, *Dunaliella salina*, *Micromonas pusilla* CCMP1545, *Micromonas* sp. RCC299, *Ostreococcus lucimarinus*, *Ostreococcus* sp. RCC809, *Ostreococcus tauri* RCC4221, *Volvox carteri*; two Pelagophyta: *Aureococcus anophagefferens* clone 1984, *Pelagophyceae* sp. CCMP2097; two Cryptophyta: *Cryptophyceae* sp. CCMP2293, *Guillardia theta* CCMP2712; three Rhodophyta: *Cyanidioschyzon merolae*, *Galdieria sulphuraria* 074W, *Porphyra umbilicalis*; four Haptophyta: *Emiliania huxleyi* CCMP1516, *Pavlovales* sp. CCMP2436, *Phaeocystis antarctica* CCMP1374, *Phaeocystis globosa* Pg-G; four Bacillariophyta: *Fragilariopsis cylindrus* CCMP 1102, *Phaeodactylum tricornutum* CCAP 1055/1, *Pseudo-nitzschia* multiseries CLN-47, *Thalassiosira pseudonana* CCMP 1335; three Ochrophyta: *Ectocarpus siliculosus* Ec 32, *Nannochloropsis oceanica* CCMP1779, *Ochromonadaceae* sp. CCMP2298; and one Chlorarachniophyta: *Bigelowiella natans* CCMP2755. In addition, other species genomes were screened for comparison from the cyanobacterium *Nostoc punctiforme* PCC 73102, human *Homo sapiens*, land plant *Arabidopsis thaliana*, yeast *Saccharomyces cerevisiae*, and the thraustochytrid *Schizochytrium aggregatum* ATCC 28209. Finally sequences from ten bacteria and one archeae were used. Genome details and individual ID numbers or accession numbers for the sequences used are provided in Supplementary Information and Table S1.

Genomes were screened by BLAST using *C. reinhardtii* and *S. cerevisiae* CDF protein sequences. In addition, genomes were screened for the presence of the Pfam domain PF01545. All sequences were then manually examined by sequence alignment to remove any false positive sequences that had been mis-annotated. Amino acid multiple sequence alignments were performed using Clustal Omega (Madeira et al. 2019). For visual analysis sequence alignments were shaded using BoxShade v.3.21 and manually annotated. Prediction of transmembrane domains (TMD) was performed by hydropathy analysis using TMHMM v.2.0 (Krogh et al. 2001), and was performed by consensus topology prediction using TOPCONS (Tsirigos et al. 2015). His-rich loop presence was determined by manual analysis by identifying the presence of three or more histidine residues within the putative cytosolic loop region between TMD 4 and 5. Conserved residues within TMD2 and TMD5 of Group 1 (Zn-CDF) and Group 5 sequences were identified using WebLogo v.2.8.2 (Crooks et al. 2004).

Phylogenetic relationships at the amino acid level were performed using full length sequences, essentially as described previously (Emery et al. 2012; Pittman and Hirschi 2016). Sequence alignment outputs from Clustal Omega were used to generate maximum likelihood phylogenies under the WAG+F model of amino acid substitution with Γ-distributed rates across sites, as implemented in RAxML v.7.1, with tree confidence determined from 1000 replications using the fast bootstrap method (Stamatakis 2006; Stamatakis et al. 2008). The trees were viewed using FigTree v.1.3 (http://tree.bio.ed.ac.uk/software/figtree).

The relationship between sequences was further quantified using Cluster Analysis of Sequences (CLANS) (Frickey and Lupas 2004), which was used to conduct an all-against-all BLAST of unaligned sequences. Pairwise similarities between protein sequences were displayed in a 2-dimensional cluster plot using CLANS Jar from the MPI Bioinformatics Toolkit (Zimmermann et al. 2018). The P value was set to 1 × 10^−5^ and the analysis was run for 50,000 iterations.

### Microalgae cultivation and RNA isolation

*C. reinhardtii* (CC125) was obtained from the Chlamydomonas Resource Center, USA, and was grown photo-heterotrophically in batch culture in Tris–acetate–phosphate (TAP) medium buffered at pH 7.0, as described previously (Harris 1989), in 200 mL glass flasks on an orbital shaker rotating at 2 Hz, at 25°C under cool-white fluorescent lights (150 μmol m^−2^ s^−1^) with a 16-h:8-h light:dark regime. Cultures were inoculated with the same starting cell density as determined by cell counting to give an initial cell count of ~65 × 10^3^ cells mL^−1^. RNA was isolated from exponential growing *C. reinhardtii* CC125 cells using Trizol reagent (Life Technologies) and further purified by phenol/chloroform extraction and precipitation with isopropanol.

### MTP full length cDNA cloning

*C. reinhardtii* genomic sequence and gene model information was obtained from Phytozome v.12 using v.5.5 of the *C. reinhardtii* genome annotations and gene-specific oligonucleotide primers incorporating restriction enzyme sites (Table S2) were designed to amplify the full-length *CrMTP1*, *CrMTP2*, and *CrMTP3* cDNA sequences by RT-PCR from *C. reinhardtii* CC125 RNA. First-strand cDNA was generated from 1 μg of DNase-treated RNA using Superscript III reverse transcriptase (Life Technologies) and an oligo(dT) primer, then KAPA HiFi DNA polymerase (Kapa Biosystems) and gene-specific primers were used for each PCR amplification, with an annealing temperature of 60°C and 35 amplification cycles. Following amplification, the PCR products were cloned into pGEM-T Easy plasmid (Promega) for propagation and sequencing (GATC Biotech) to confirm sequence fidelity. The yeast expression plasmid piUGpd was used to allow expression of each cDNA under the control of the constitutive yeast GAPDH promoter and selection of the *URA3* gene (Nathan et al. 1999). *CrMTP1* cDNA was sub-cloned into the XbaI and SacI sites of piUGpd, while *CrMTP2* and *CrMTP3* cDNA were sub-cloned into the BamHI and XbaI sites of piUGpd. *CrMTP4* in piUGpd was generated previously (Ibuot et al. 2017).

### Quantitative gene expression analysis

*C. reinhardtii* CC125 was cultured in TAP medium for three days then cells were harvested and frozen in liquid N_2_. RNA was isolated and cDNA was produced as described above. Expression for each MTP gene was determined by quantitative real-time PCR (qPCR) using short amplicon primers that were designed to be specific to each gene (Table S3). For qPCR a SYBR Green core qPCR kit (Eurogentec) and a StepOnePlus machine (ThermoFisher) using the SYBR Green detection program was used. Transcript abundance for each *MTP* was normalised to *CBLP* transcript abundance. Reactions were run in triplicate and qPCR analysis was performed as described previously (Bajhaiya et al. 2016).

### Yeast heterologous expression and metal tolerance assays

Yeast (*S. cerevisiae*) strains *pmr1* (*MATa; his3*∆1; *leu2*∆*0*; *met15*∆*0*; *ura3*∆*0*; *pmr1*::*kanMX4*) (Euroscarf, Frankfurt, Germany), *zrc1 cot1* (*MATa; his3*∆1; *leu2*∆*0*; *met15*∆*0*; *ura3*∆*0*; *zrc1*::*natMX*; *cot1*::*kanMX4*) (Drager et al. 2004) (kindly provided by Ute Krämer) and the corresponding wild type strain BY4741 (*MATa; his3*∆1; *leu2*∆*0*; *met15*∆*0*; *ura3*∆*0*) (Euroscarf) were each transformed with each *MTP-*piUGpd plasmid or empty piUGpd plasmid using the lithium acetate-polyethylene glycol method. Colonies were selected by growth at 30°C in synthetic defined medium minus uracil (SD −Ura) as described previously (Pittman et al. 2004). Expression of each *MTP* cDNA in yeast was confirmed by RT-PCR using the appropriate *MTP* qPCR primer pair (Table S3) and RNA extracted from yeast using Trizol reagent, then RT-PCR was performed as described above. PCR products were examined on a 1% agarose gel stained with SafeView (NBS Biologicals).

Metal tolerance assays were performed essentially as described previously (Pittman et al. 2009), on solid SD −Ura medium with or without 2 mM ZnCl_2_, 5 mM MnCl_2_ or 0.8 mM CoCl_2_ metal salts. These concentrations were chosen as the most appropriate to restrict growth of the Zn- and Co-sensitive *zrt1 cot1* strain and the Mn-sensitive *pmr1* strain but still allow the wild type BY4741 to grow. For growth in liquid SD −Ura medium, yeast strains were grown in medium with or without 10 mM or 20 mM CoCl_2_, MnCl_2_, or ZnCl_2_ metal salts.

Cell density was determined by absorbance measurement at 600 nm while yeast biomass was determined by weighing a cell pellet from a set volume of culture after it was dried at 60°C for 24 h. Internal metal content in yeast grown in liquid SD −Ura medium containing 20 mM ZnCl_2_ or 20 mM MnCl_2_, respectively, was determined by inductively coupled plasma atomic emission spectroscopy (ICP-AES) essentially as described previously using a Perkin-Elmer Optima 5300 (Webster et al. 2011). It was estimated using Visual Minteq v.3.0 metal speciation modelling that the free ion concentrations available to the cells was 10.4 mM Zn^2+^ and 12.6 mM Mn^2+^. Cells were centrifuged for 10 min at 3000 g then resuspended in 10 mL of 10 mM EDTA for 5 min, then re-centrifuged and washed with 15 mL Milli-Q water. Cell pellets were dried at 60 °C for 24 h and then digested in 0.5 mL of ultrapure concentrated nitric acid at 70 °C for 2 h. Samples were diluted in Milli-Q water to 2% (v/v) concentration of acid and analysed by ICP-AES. All samples were calibrated using a matrix-matched serial dilution of Specpure multi-element plasma standard solution 4 (Alfa Aesar) set by linear regression.

## Results

### Phylogenetic analysis of algal CDF protein sequences

Previous phylogenetic analyses of CDF sequences are inconsistent in terms of clade structure and nomenclature of groups, with proposed numbers of CDF sub-groups ranging from 3 to 18 groups (Migeon et al. 2010; Cubillas et al. 2013; Gustin et al. 2011; Montanini et al. 2007). Montanini et al. (2007) divided the CDF family into three major groups based on substrate specificity, referred to as Zn-CDF, Fe/Zn-CDF and Mn-CDF, while Gustin et al. (2011) further sub-divided the higher plant CDF members into seven groups. Cubillas et al. (2013) identified 18 CDF groups that were differentiated mostly depending on whether they contained eukaryote, bacteria or archaea derived sequences. Here we performed a phylogenetic analysis that focussed on available algal genomes, including from one charophyte, 13 chlorophytes, two pelagophytes, two cryptophytes, three rhodophytes, four haptophytes, four bacillariophytes, three ochrophytes and one chlorarachniophyte (Table 1). A representative ‘algal-like’ thraustochytrid (*S. aggregatum*) was also included. First, each genome was analysed by a combination of BLAST and Pfam signature searching to collate CDF sequences. Each algal genome examined had at least one CDF sequence and the numbers of CDF transporter genes ranged from one in the red alga *G. sulphuraria* to 14 genes in the haptophyte bloom microalga *P. antarctica* (Table 1). The exact numbers of CDF genes within the more recently annotated genomes should be considered with caution since incorrect annotation and genome assembly errors can lead to genes being missed.

**Table 1.**
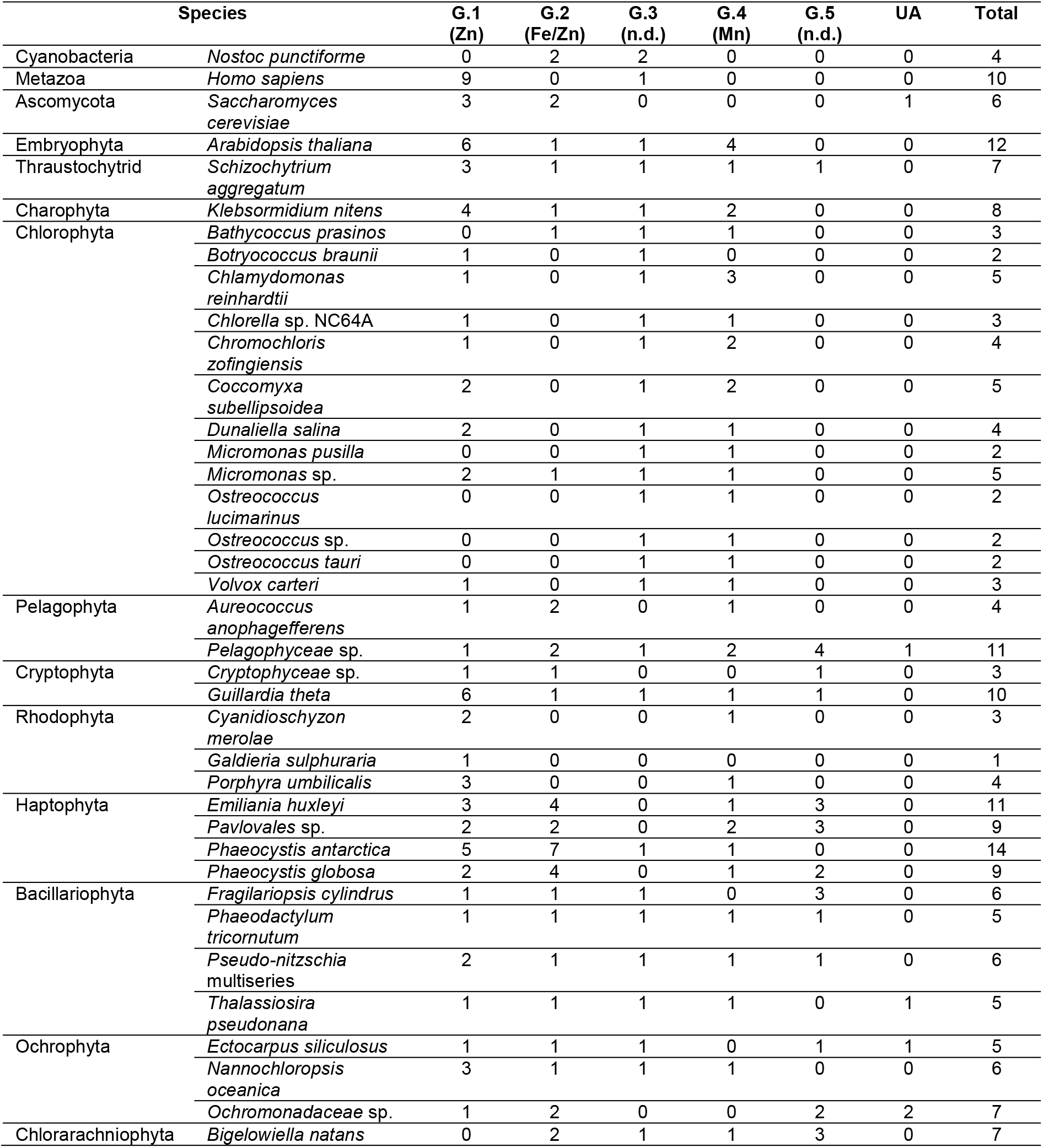
Total number and distribution of CDF transporter genes from each analysed genome into five phylogenetic groups: Group 1 (Zn-CDF), Group 2 (Fe/Zn-CDF), Group 3 (undefined CDF), Group 4 (Mn-CDF), Group 5 (undefined CDF). Gene numbers were determined from the phylogenetic analysis (Fig. 1) and supported by the cluster analysis (Fig. 2). Six sequences were classed as unassigned (UA) including five sequences where the evidence for Group 2 assignment was not supported by cluster analysis.

To generate the phylogenetic tree, the algal sequences were aligned with representative sequences from bacteria (*Escherichia coli*, *Bacillus subtilis*, *Bradyrhizobium diazoefficiens*, *Cupriavidus metallidurans*, *Rhizobium etli*, *Sinorhizobium meliloti*, *Streptococcus pneumoniae*, *Thermus thermophiles*), archaea (*Methanosarcina acetivorans*), cyanobacteria (*N. punctiforme*), animals (*H. sapiens*), fungi (*S. cerevisiae*), and land plants (*A. thaliana*), which had all been used in previous phylogenetic analyses and in most cases have known substrate specificity (Table S1). The constructed phylogenetic tree contained 226 sequences, including 15 prokaryotic, 10 human, 12 plant, 6 yeast, 7 thraustochytrid and 176 algal sequences, of which 42 were chlorophyte sequences. Despite the dominance of many new algal sequences, the tree structure was consistent with previous CDF trees (Gustin et al. 2011; Montanini et al. 2007). It possessed distinct groups containing many of the known Zn^2+^, Fe^2+^ and Mn^2+^ transporting CDF proteins characterised in human, yeast, *A. thaliana* and bacteria, which were recognisable as the previously named Zn-CDF, Fe/Zn-CDF and Mn-CDF groups (Fig. 1). Group 1 (Zn-CDF) could have been considered as two or three separate groups, including a bacterial-specific clade (shown in dark blue in Fig. 1) and two eukaryotic groups (an AtMTP1 − AtMTP4 containing group, and an AtMTP5 and AtMTP12 containing group), which all include CDF proteins with experimentally confirmed Zn^2+^ transport activity (e.g. Arrivault et al. 2006; Fujiwara et al. 2015; Chao and Fu 2004; Liuzzi and Cousins 2004; Kobae et al. 2004; MacDiarmid et al. 2002), indicating functional conservation across the whole Group 1. All taxonomic classes of algae possessed a Group 1 Zn-CDF, apart from the Chlorarachniophyta *B. natans*, although 5 of the 13 chlorophytes, including all three *Ostreococcus* spp. appear to lack a Zn-CDF (Table 1).

**Fig. 1.**
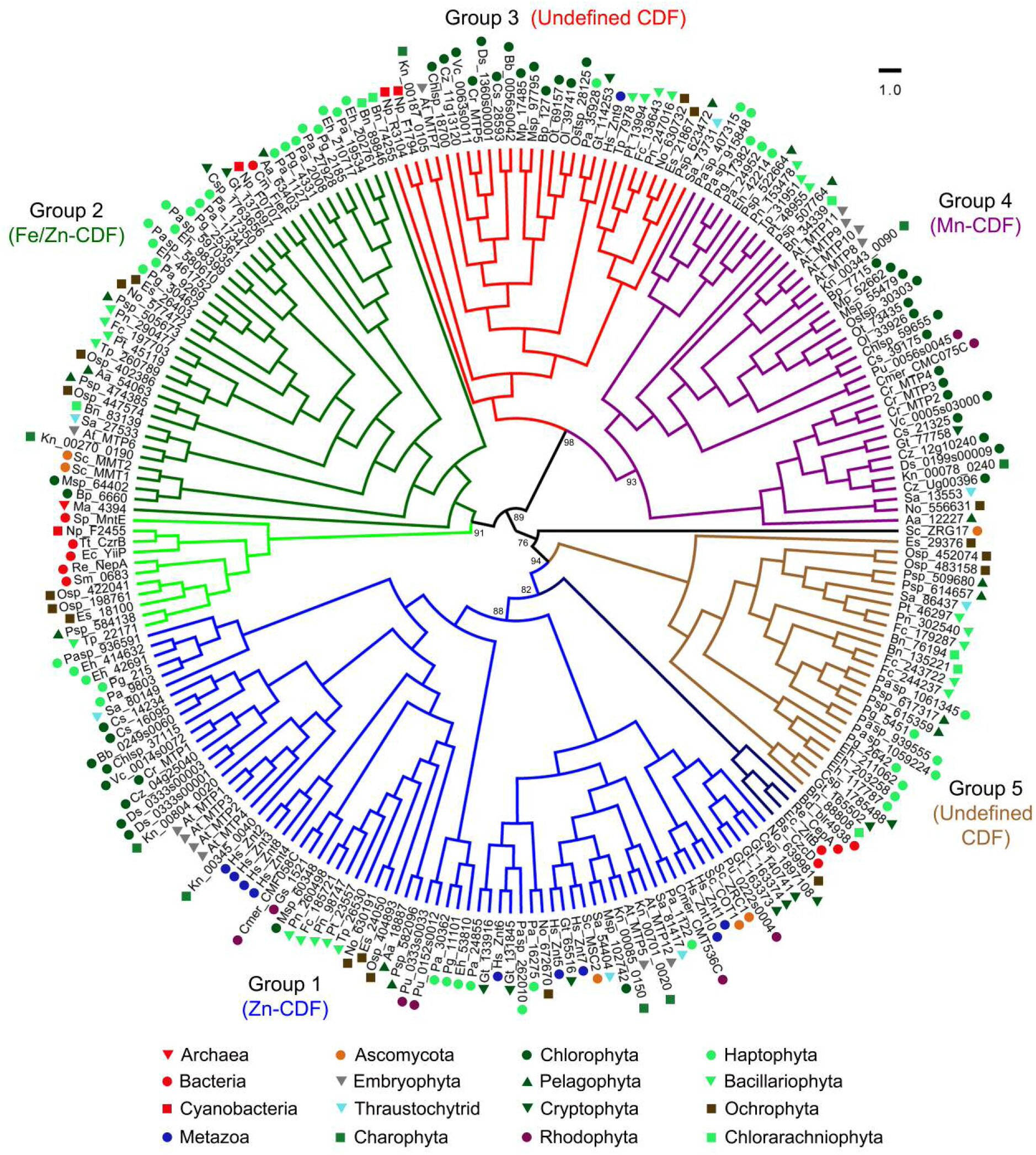
Phylogenetic analysis of CDF transporter family proteins from various selected algal taxa in comparison to prokaryotic, animal (*H. sapiens*), land plant (*A. thaliana*) and yeast (*S. cerevisiae*) proteins. The genome identifier or accession numbers of all sequences used are provided in Table S1. Coloured symbols indicate the major taxonomic domains of the species where the protein sequence is derived. Five major groups are highlighted. The tree was constructed using the maximum likelihood method and derived from alignments of full length amino acid sequence. A consensus tree following 1000 bootstrap replications is shown, with bootstrap percentage values indicated at the nodes of major branches. The branch length scale bar indicates the evolutionary distance of one amino acid substitution per site.

Group 2 (Fe/Zn-CDF), which includes Fe^2+^ and/or Zn^2+^ transporting proteins such as yeast ScMMT1 and ScMMT2, and *C. metallidurans* CmFieF, is rich in algal CDF sequences, including a large number of haptophyte, bacillariophyte, and ochrophyte sequences, but includes only two chlorophyte and no rhodophyte sequences (Fig. 1). A predominantly bacterial sub-group also containing five algal sequences (shown in light green in Fig. 1) might be considered as a separate group since it contains CDF proteins with known Mn^2+^, Ni^2+^ and Co^2+^ transport activities (Cubillas et al. 2013), rather than Fe^2+^/Zn^2+^. Group 3 and Group 4 (Mn-CDF) are considered as two separate groups rather than a single group. Group 4 contains many known plant Mn^2+^ transporting CDFs, such as AtMTP11 (Delhaize et al. 2007; Peiter et al. 2007), and is also enriched in chlorophyte and haptophyte sequences. Group 3 is also rich in sequences from all chlorophyte species, and includes CDF proteins such as AtMTP7 and HsZnt9 that have no currently confirmed substrate (Fig. 1). Although indirect evidence indicates Zn^2+^ transport activity by HsZnt9 (Perez et al. 2017), Group 3 is referred to as a separate undefined CDF group. Finally, the tree displays a fifth algal-specific group, which is also referred to as undefined. Group 5 sequences are missing in all of the examined higher plant, charophyte and chlorophyte genomes (Fig. S2).

In order to provide validation for the group structure proposed for the CDF tree shown in Fig. 1, a phenetic sequence clustering method was performed using CLANS (Frickey and Lupas 2004). Nearly all of the sequences fell into five clusters, which completely matched the Groups 1 to 5 determined by the phylogenetic tree (Fig. 2). In particular, the analysis fully validated the separation of Group 3 and 4 into independent groups, which were both very tightly clustered. The Group 2 sequences were fairly tightly clustered and indicated partly that this group should not be further divided (shown by light green and dark green symbols overlapping in Fig. 2), although six of the sequences within this sub-group, including five algal proteins from *Pelagophyceae* sp., *T. pseudonana*, *E. siliculosus* and *Ochromonadaceae* sp., and the bacterial Sm_0683 protein from *S. meliloti* did not cluster with the other Group 2 sequences, and are instead classified as ‘unassigned’ (Table 1). Group 1 and 5 sequences also formed clearly distinct clusters although these were not as tightly clustered as the other groups, and there was some overlap between four of the Group 5 sequences with Group 1. Overall, this CLANS analysis validated the five-group organisation for these CDF sequences.

**Fig. 2.**
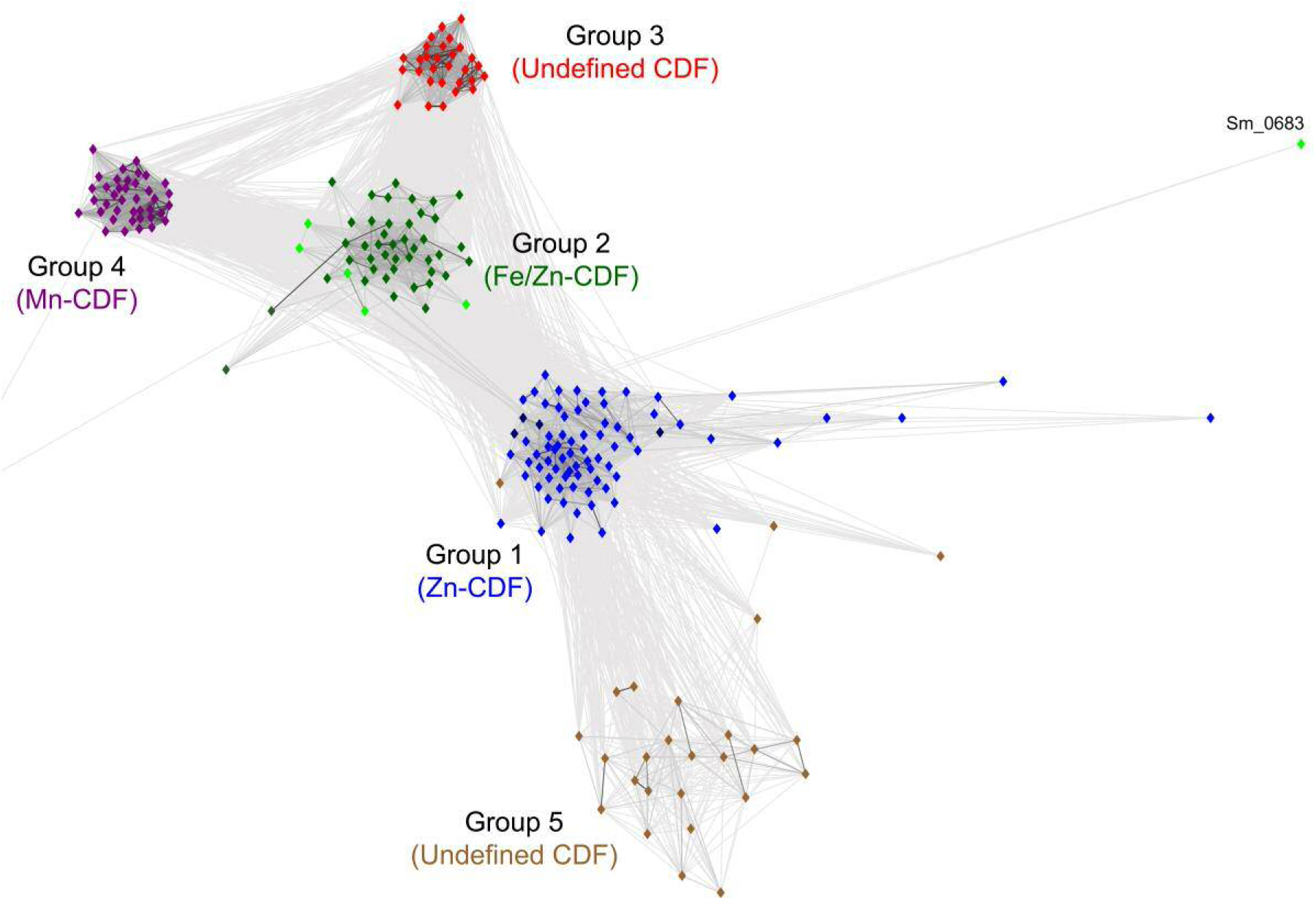
A two-dimensional cluster analysis of sequences representing the phenetic sequence relationship of CDF transporter family proteins from various selected algal taxa in comparison to prokaryotic, animal (*H. sapiens*), land plant (*A. thaliana*) and yeast (*S. cerevisiae*) proteins. Each symbol represents a full length amino acid sequence of an individual CDF protein and is coloured on the basis of the phylogenetic groups shown in Fig.1. Grey lines represent protein connections with reciprocal BLAST hits at P < 10^−5^. Six proteins are not shown (five Group 2 members: Psp_584138, Tp_22171, Es_18100, Osp_198761, Osp_422041, and unassigned ScZRG17) as they did not cluster with any of the other sequences and are on the periphery of the plot off the scale.

### Structural features of Zn-CDF related sequences from algae

Conserved amino acid residues within TMD2 and 5 have been demonstrated to be critical for metal ion binding, in particular, two residues from TMD2 and two from TMD5 form a metal binding site referred to as ‘binding site A’ (Kolaj-Robin et al. 2015; Barber-Zucker et al. 2017). The presence of residues HxxxD-HxxxD within the TMD2 and TMD5 regions, respectively, was conserved for the algal-dominated Group 1 proteins examined here (Fig. 3), which is consistent with previous analysis of Zn-CDF proteins (Montanini et al. 2007). In contrast, the Group 5 proteins displayed a binding site A motif that was predominantly A/SxxxD-HxxxD, such that Ala or Ser replaced the conserved His (Fig. 3). Only 4 of the 26 Group 5 proteins had the HxxxD-HxxxD motif that was highly conserved within Group 1 proteins. Indeed, these were the four Group 5 proteins that partially overlapped with the Group 1 cluster (Fig. 2).

**Fig. 3.**
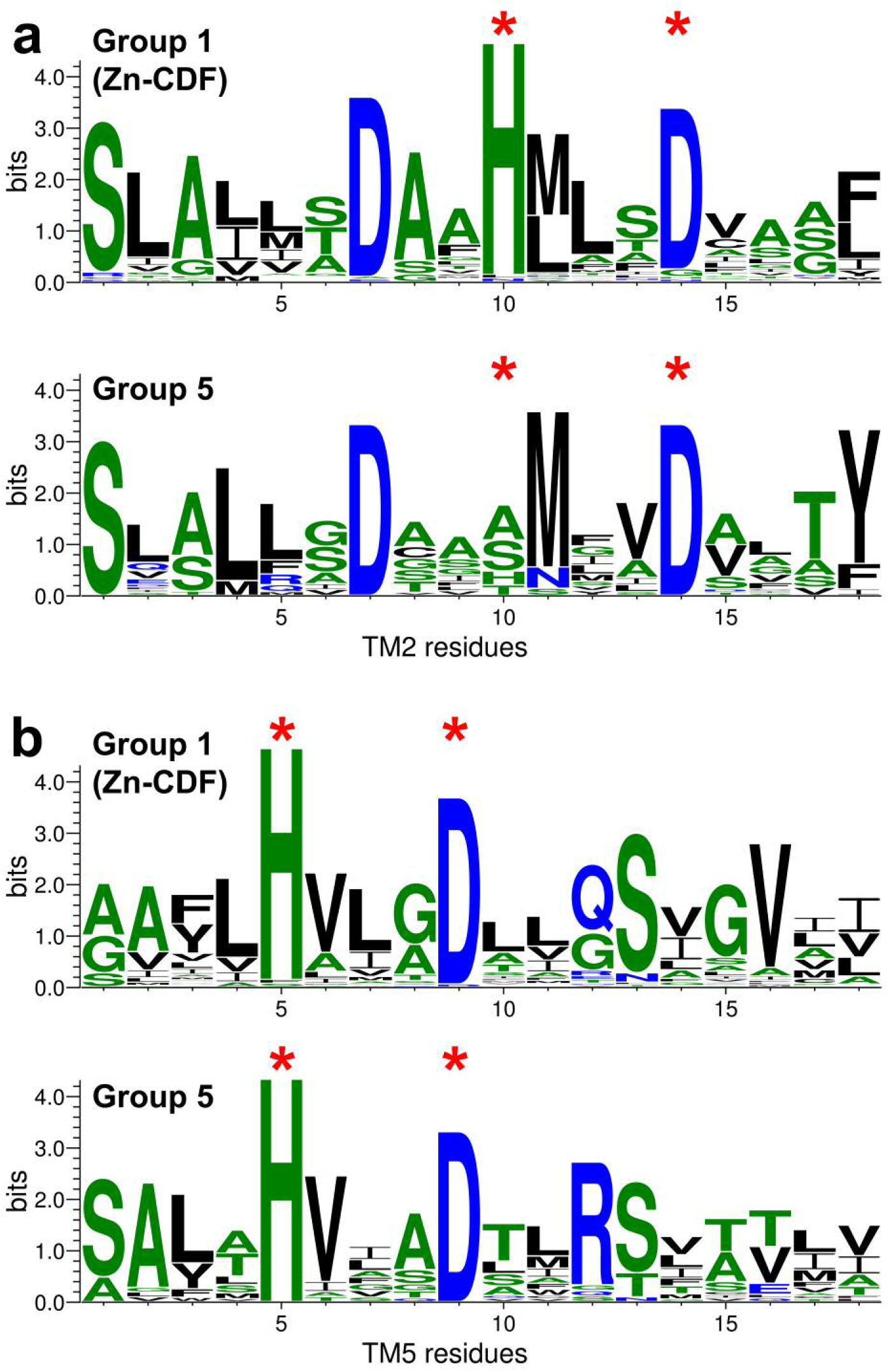
Comparison of conserved CDF residues within transmembrane (TM) domain 2 (**a**) and TM domain 5 (**b**) for Group 1 (Zn-CDF) and Group 5 proteins. Sequence logo representations of 18 residues within each TM domain determined for all of the Group 1 sequences and all of the Group 5 sequences. Asterisks indicate the positions of conserved active site ‘A’ residues.

Most of the Zn^2+^-transporting CDF proteins within Group 1 possess a His-rich cytosolic loop between TMD4 and 5, and this loop may play a role in metal selectivity, as some Zn-CDF transporters are specific for Zn^2+^ while others can also transport metals including Co^2+^ (Kawachi et al. 2012; Podar et al. 2012). However, the presence of a His-rich sequence is not universal, and while most the Zn-CDF proteins examined here possess an obvious His-rich loop region, including most of the chlorophyte and bacillariophyte sequences, a few do not, including some of the human Znt proteins, *A. thaliana* AtMTP5, and AtMTP5-related algal sequences (Fig. 4a). Another notable feature of many of the algal Zn-CDF sequences is that the His-rich cytosolic loop is often significantly longer than the equivalent loop region present within CDF proteins such as from *Arabidopsis*, human and yeast (Fig. 4b). For example, the CrMTP1 His-rich loop is over 300 amino acids in length while the AtMTP1 loop just over 50 amino acids in length (Fig. S3). However, some algal CDF proteins, such as from *Micromonas* sp. and *G. theta* have very short His-stretch regions of less than 10 amino acids (Fig. 4b).

**Fig. 4.**
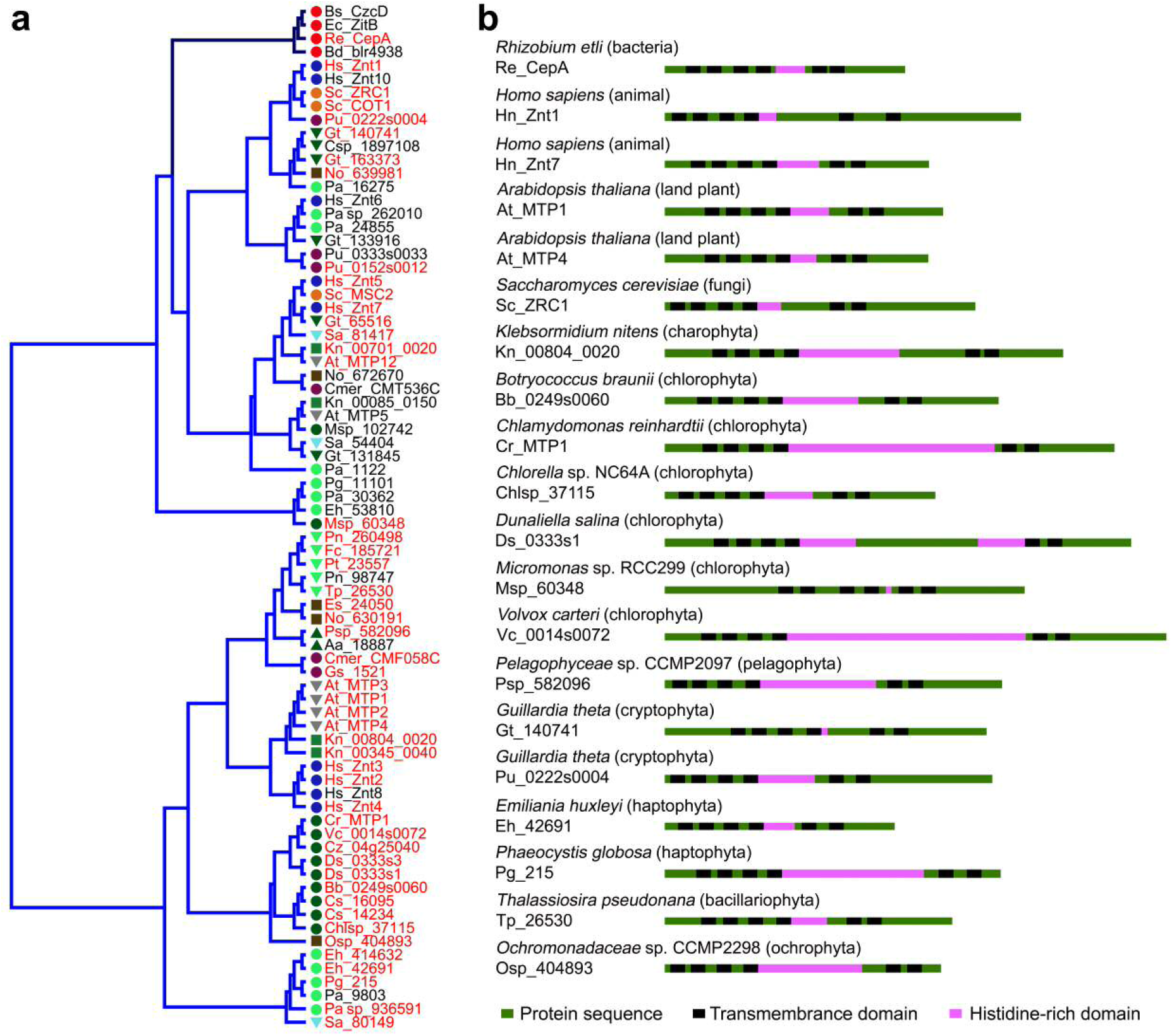
Analysis of Group 1 Zn-CDF proteins. (**a**) Phylogenetic analysis of Zn-CDF family proteins from various selected species. The coloured symbols indicating the major taxonomic domains of each species are as defined in Fig. 1. Proteins highlighted in red possess an obvious histidine-rich domain. (**b**) Topology models of selected Group 1 Zn-CDF proteins indicating the predicted transmembrane (TM) spans and the histidine-rich domain within the major cytosolic loop between TM4 and TM5.

### Functional characterisation of C. reinhardtii CDF proteins

*C. reinhardtii* has five CDF proteins, CrMTP1 within Group 1 (Zn-CDF), CrMTP2, CrMTP3 and CrMTP4, within Group 4 (Mn-CDF), and CrMTP5 within the undefined Group 3 (Fig. S2). To determine whether all genes are expressed within the organism, RT-PCR was performed using RNA extracted from exponential growth stage cells grown under standard conditions. Transcripts were detected for all five genes with *CrMTP1*, *CrMTP2* and *CrMTP5* showing an equivalent level of expression (between approximately 0.2 − 0.3% of *CBLP* expression), while *CrMTP3* and *CrMTP4* had low but detectable levels of expression that was significantly reduced (P < 0.05) by approximately 10 − 20-fold compared to that of *CrMTP1*, *CrMTP2* and *CrMTP5* (Fig. 5). Growth of *C. reinhardtii* in excess concentrations of metals such as Cd, Co, Fe, Mn and Zn had no significant effect on the expression level of any of the genes (data not shown) indicating that none of these CrMTP genes are transcriptionally regulated by excess metal status.

**Fig. 5.**
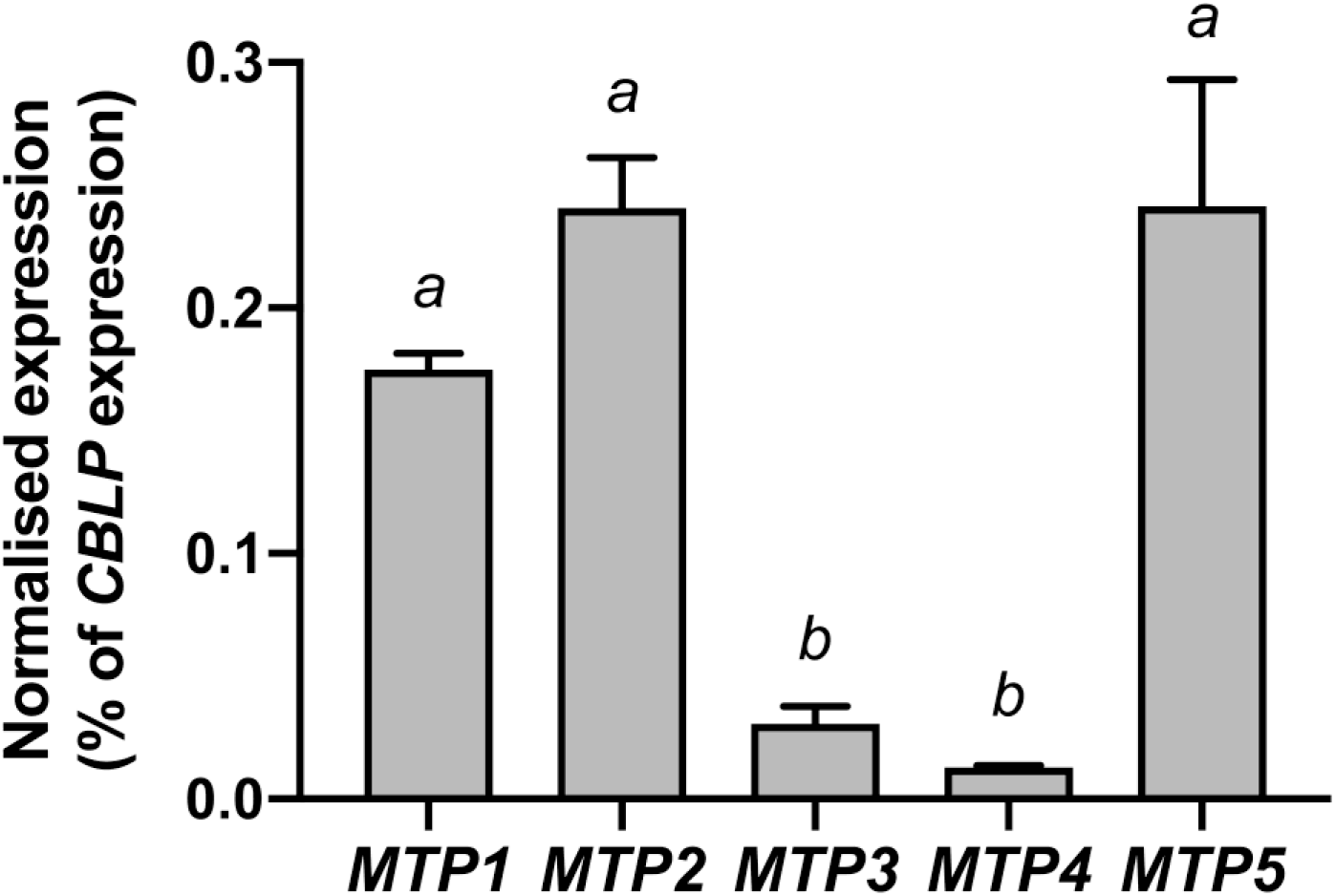
Expression the CrMTP family genes in *C. reinhardtii* cells grown in standard TAP medium at exponential growth stage. Expression of the selected mRNA transcripts as determined by real-time PCR was calculated as a percentage of *CBLP* expression. Data points are means (±SEM) calculated from biological triplicates. Bars sharing the same lowercase letter indicate no significant difference between treatments (P > 0.05) as determined by one-way ANOVA and Tukey post-hoc test.

CrMTP4 has previously been confirmed to have Mn^2+^ transport activity and the ability to transport Cd^2+^ (Ibuot et al. 2017), but the substrate specificity of the other *C. reinhardtii* CDF proteins is unknown. The full-length cDNA for each gene was functionally characterised by heterologous expression in yeast (Fig. 6), however, the expression of CrMTP5 was unsuccessful and therefore no functional information could be obtained for this protein. In contrast, CrMTP1, CrMTP2, CrMTP3 and CrMTP4 were all successfully expressed in a Zn- and Co-sensitive yeast strain *zrc1 cot1* (lacking the yeast vacuolar Zn^2+^ and Co^2+^ transporters ScZRC1 and ScCOT1, respectively), and a Mn-sensitive yeast strain *pmr1* (lacking the Mn^2+^ ATPase transporter ScPMR1) (Fig. 6b). All cDNAs were functional as indicated by their ability to suppress metal sensitive phenotypes (Fig. 6a). CrMTP4 was used as a positive control as a confirmed Mn-CDF, and as seen with other higher plant Mn-CDF proteins, it was unable to rescue the growth of *zrc1 cot1* yeast on Zn or Co media. In contrast, CrMTP1 could strongly suppress both the Zn and Co sensitivity of *zrc1 cot1* but was unable to suppress the Mn sensitivity of *pmr1*, indicating that CrMTP1 is a Zn^2+^ and Co^2+^ transporter. CrMTP2 showed the identical characteristics of CrMTP4, indicating that it is a specific Mn^2+^ transporter (Fig. 6a). CrMTP3 was interesting as it could strongly suppress the Mn sensitivity of *pmr1* but could also partially suppress the Zn and Co sensitivity of *zrc1 cot1*, although not as strongly as CrMTP1. This suggested that CrMTP3 has broad substrate specificity for all three divalent cations tested. To date, none of the Mn-CDF proteins have been found to have Zn^2+^ or Co^2+^ transport activity, and appear to be largely Mn^2+^-specific (Delhaize et al. 2007; Montanini et al. 2007; Peiter et al. 2007).

**Fig. 6.**
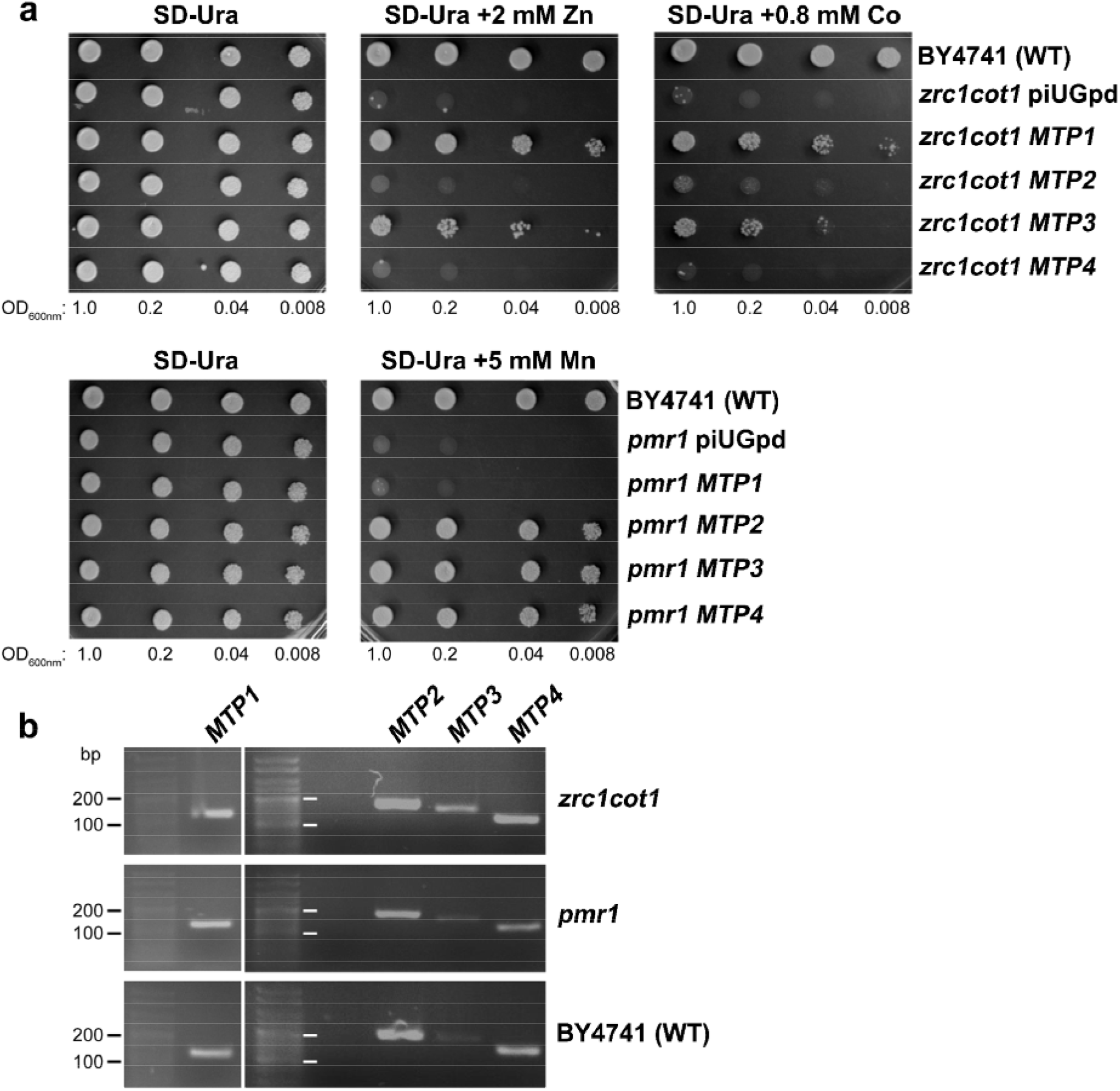
Heterologous expression of *CrMTP1*, *CrMTP2*, *CrMTP3* and *CrMTP4* in yeast strains. (**a**) Suppression of Zn, Co and Mn hypersensitivity of the Zn- and Co-sensitive *zrc1cot1* yeast mutant and the Mn-sensitive *pmr1* mutant by heterologous expression of *CrMTP* cDNA in comparison with empty vector (piUGpd) and WT (BY4741). Saturated liquid cultures of yeast strains were serially diluted to the indicated cell densities then spotted onto SD −Ura selection medium with or without added Zn, Co or Mn. Yeast growth is shown after 3 d. A representative experiment is shown. (**b**) RT-PCR of *MTP1*, *MTP2*, *MTP3* and *MTP4* expression in each yeast strain.

To further confirm the Zn, Co and Mn tolerance abilities of these proteins all cDNAs were expressed in a wild type yeast strain (BY4741) (Fig. 6b). Growth of the yeast in liquid medium was quantified in the presence of external metal addition, at concentrations that induced toxicity by inhibiting growth of wild type yeast and were relevant to metal contaminated environments (Wuana and Okieimen 2011). CrMTP1 but not CrMTP3 was able to moderately but significantly increase tolerance to Zn in the yeast strain as determined by cell density measurement (Fig. 7b) and by yeast biomass (Fig. 8a). However, there was no significant increase in yeast growth for any of the MTP-expressing strains in response to Co addition (Fig. 7c and Fig. 8b). All three CDF proteins (CrMTP2, CrMTP3, CrMTP4) that were able to strongly suppress the Mn sensitivity of *pmr1* yeast could also provide strong growth tolerance to BY4741 yeast in the presence of 20 mM Mn (Fig. 7d and Fig. 8c).

**Fig. 7.**
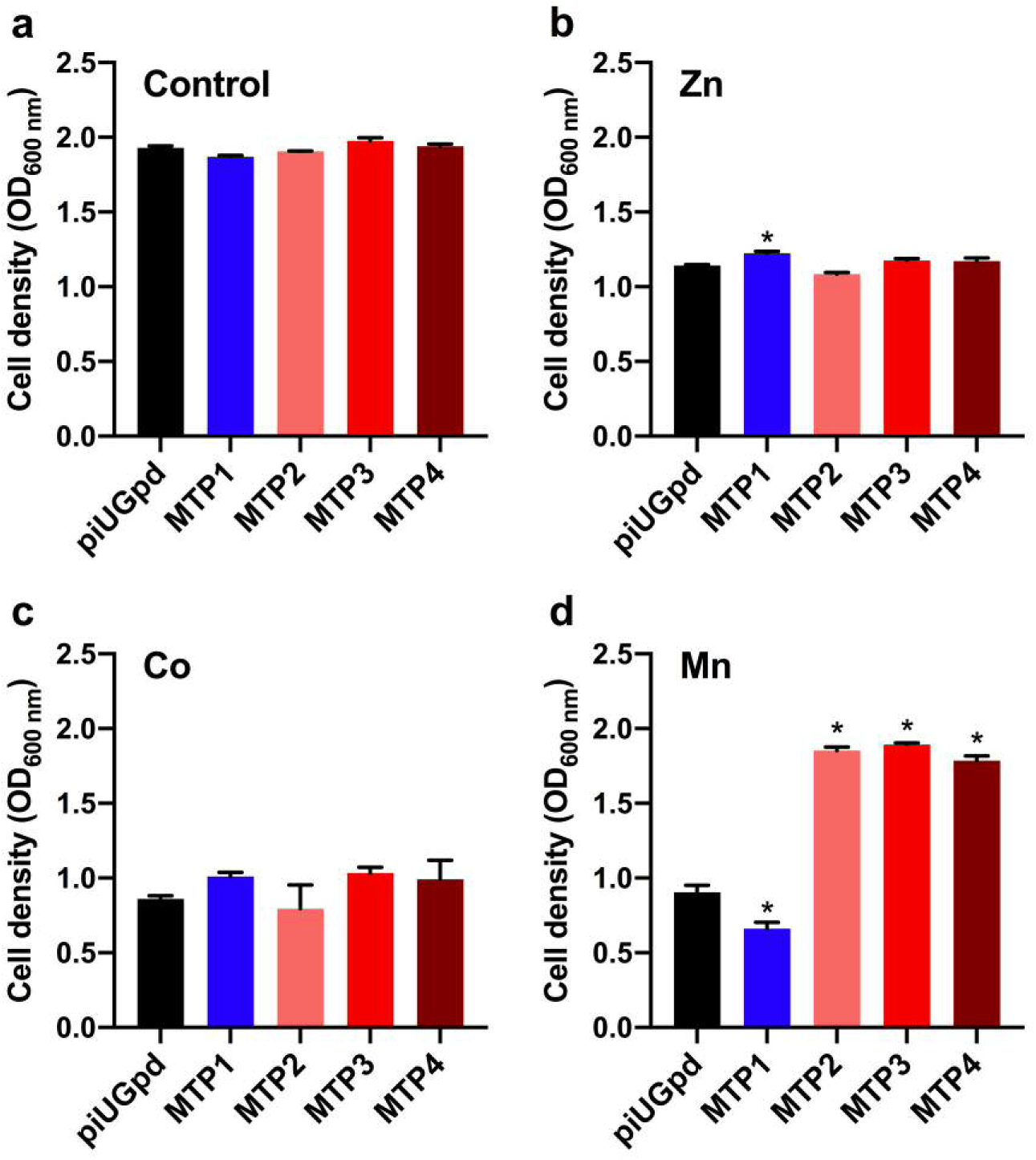
Heterologous expression of *CrMTP1*, *CrMTP2*, *CrMTP3* and *CrMTP4* in wild type (BY4741) yeast. Cell density measurement of strains expressing each MTP cDNA in comparison with empty vector (piUGpd) grown after 3 d in liquid SD −Ura medium either without metal exposure (**a**) or with 10 mM Zn (**b**), 10 mM Co (**c**), or 20 mM Mn (**d**). Data points are means (±SEM) from triplicate samples. Bars with an asterisk indicate significant difference between the empty vector control (piUGpd) strain (P < 0.05) as determined by one-way ANOVA and Tukey post-hoc test.

**Fig. 8.**
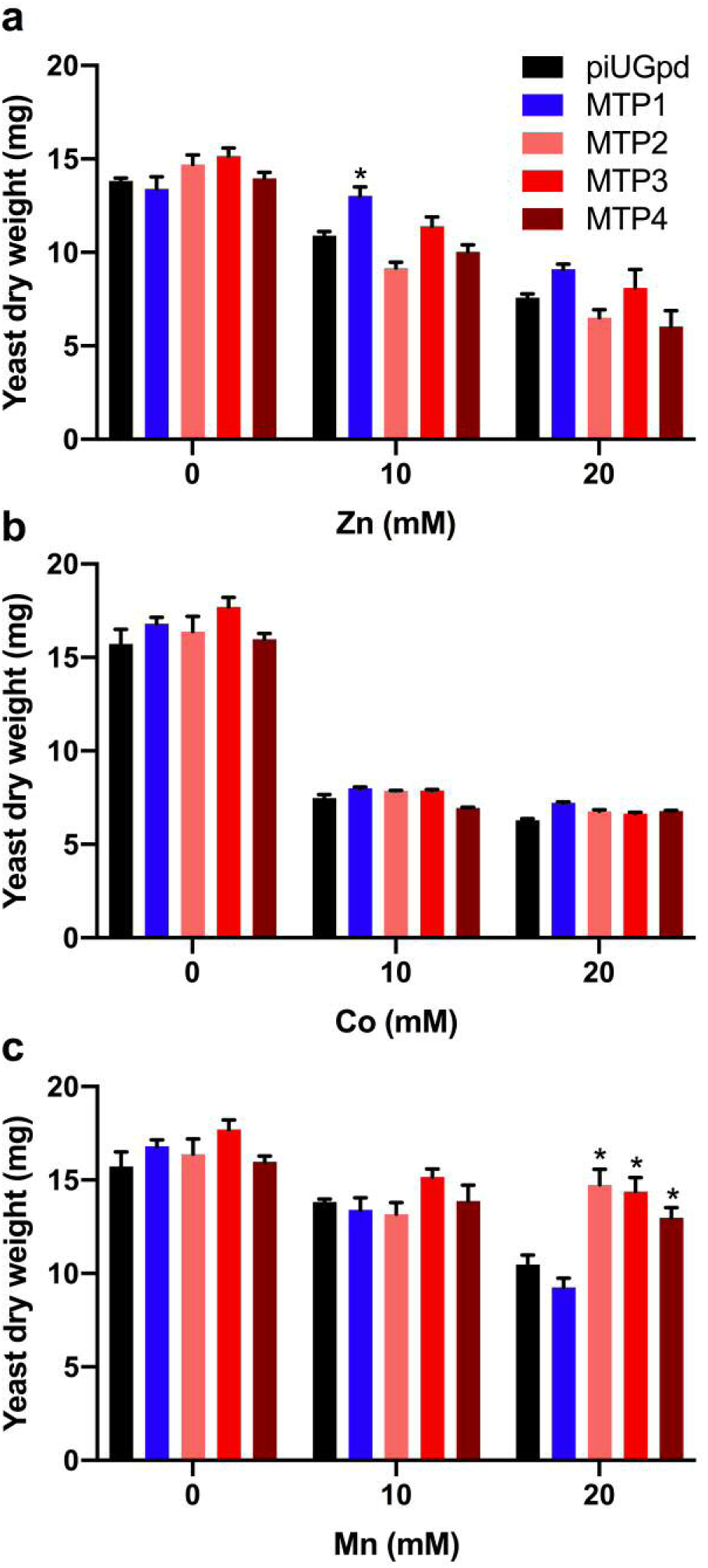
Heterologous expression of *CrMTP1*, *CrMTP2*, *CrMTP3* and *CrMTP4* in wild type (BY4741) yeast. Dry weight biomass measurement of strains expressing each *CrMTP* cDNA in comparison with empty vector (piUGpd) grown after 3 d in liquid SD −Ura medium either without metal exposure or with added Zn (**a**), Co (**b**), or Mn (**c**). Data points are means (±SEM) from triplicate samples. Bars with an asterisk indicate significant difference between the empty vector control (piUGpd) strain (P < 0.05) as determined by one-way ANOVA and Tukey post-hoc test.

### Metal accumulation into yeast expressing C. reinhardtii CDF proteins

To determine whether the expression of algal CDF proteins in yeast can provide increased metal transport into the cell, Zn and Mn concentration in yeast cell biomass was quantified by ICP-AES following growth in liquid media with added Zn or Mn. Prior to measurement, cells were washed in an EDTA solution to remove cell wall bound metals so that only internalised metals were measured. There was no significant increase in Zn content for any of the strains including for the CrMTP1 expressing yeast (Fig. 9a), whereas the yeast expressing CrMTP2, CrMTP3 and CrMTP4 showed an approximately 2-fold increase in Mn content within the cells (Fig. 9b), indicating that the Mn tolerance of these strains was potentially due to internal sequestration of Mn.

**Fig. 9.**
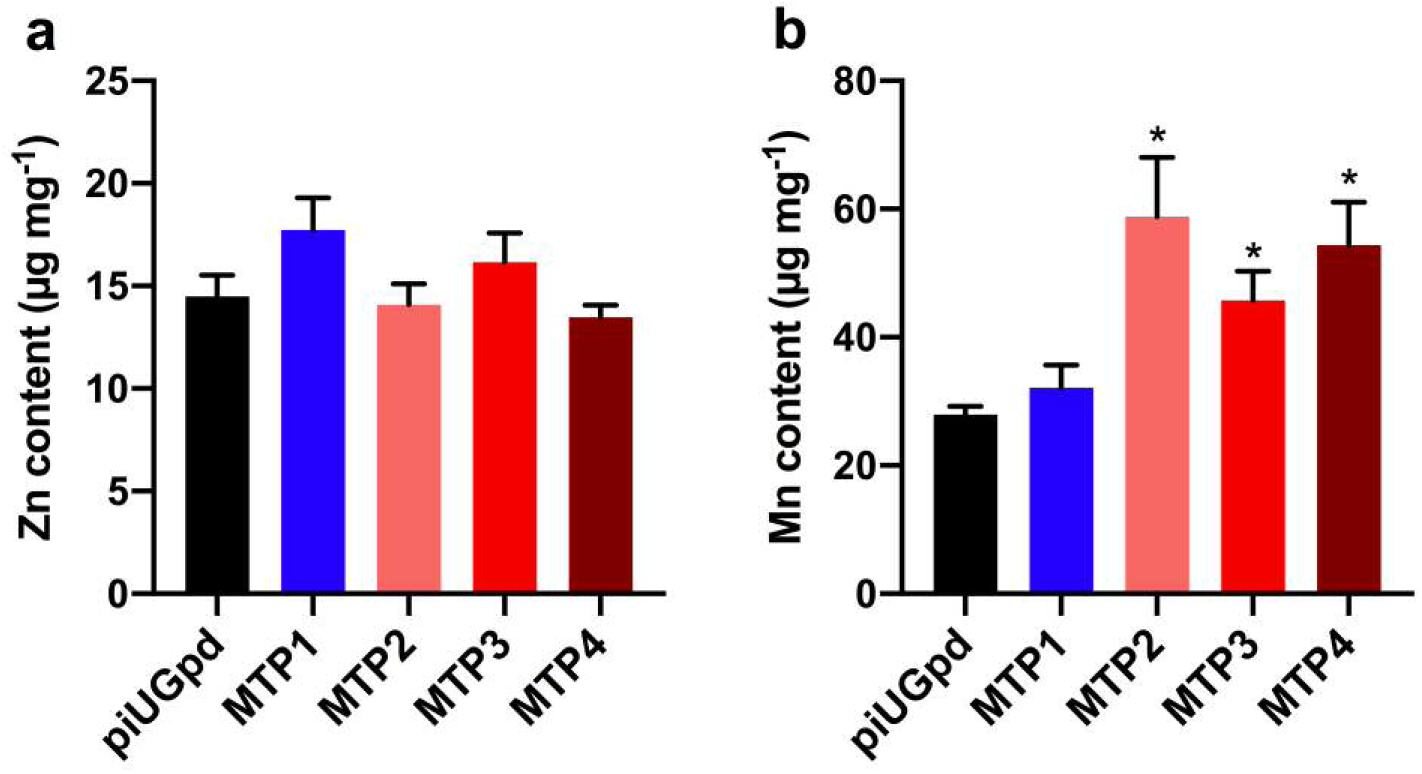
Zn and Mn uptake into wild type (BY4741) yeast expressing *CrMTP1*, *CrMTP2*, *CrMTP3* and *CrMTP4*. Metal content measurement of strains expressing each *CrMTP* cDNA in comparison with empty vector (piUGpd) grown after 3 d in liquid SD −Ura medium with added 20 mM Zn (**a**) or 20 mM Mn (**b**). Data points are means (±SEM) from triplicate samples. Bars with an asterisk indicate significant difference between the empty vector control (piUGpd) strain (P < 0.05) as determined by one-way ANOVA and Tukey post-hoc test.

## Discussion

### Conserved phylogenetic structure of algal CDF transporters

CDF metal transporters are a ubiquitous family of proteins across life, and this analysis confirms their presence across the wide range of taxonomically and physiologically diverse photosynthetic organisms that are collectively referred to as algae. CDF transporters appear to be principally involved in the transfer of various essential metal ions including Zn^2+^, Mn^2+^, Fe^2+^ and Co^2+^ into internal organelles either for delivery to specific subcellular-localised proteins that require metal co-factors or for providing metal tolerance through internal sequestration (Kolaj-Robin et al. 2015; Ricachenevsky et al. 2013). Although other classes of primary or secondary energised metal transporters can play similar roles (Blaby-Haas and Merchant 2012), CDF transporters are clearly indispensable in algae since each algal genome that has been analysed here has at least one CDF isoform. Furthermore, the algal-dominated phylogenetic analysis performed in this study was consistent with a previously determined CDF phylogenetic structure, whereby a small number of major groups can be differentiated on the basis of Zn^2+^, Fe^2+^/Zn^2+^ and Mn^2+^ substrate specificity, plus undefined substrate specificity (Montanini et al. 2007). The fact that this phylogenetic structure is conserved in many algal species suggests that it was determined early in the evolution of photosynthetic organisms. In particular, the Zn-CDF and Mn-CDF groups are the most conserved across the algal genomes examined here. Nevertheless, some algal species have lost particular CDF types; for example, all three *Ostreococcus* spp. lack Group 1 (Zn-CDF) and Group 2 (Fe/Zn-CDF) genes, and in fact most chlorophytes lack Group 2 (Fe/Zn-CDF) genes, while all three red algae examined (*C. merolae*, *G. sulphuraria* and *P. umbilicalis*) lack Group 2 (Fe/Zn-CDF) and the undefined Group 3 genes. The functional relevance of this is unclear but may indicate that functional redundancy with other transporter classes (e.g. Zn^2+^ or Mn^2+^ P-type ATPases; Fe^2+^ or Mn^2+^ transporting VIT-like proteins) is not essential and such redundancy has therefore not been selected in many of these organisms.

### A novel algal-specific phylogenetic group with distinct ion binding site motif

We have also proposed the presence of a fifth phylogenetic group (Group 5) that is most closely related to Group 1 (Zn-CDF) but appears to be distinct. This group is almost exclusively algal-specific apart from a *S. aggregatum* isoform, but includes no chlorophyte or rhodophyte sequences. The difference in sequence for the TMD2 half of the metal binding site A motif of the majority of the Group 5 proteins further demonstrates a divergence with the other groups. This TMD2 motif is typically HxxxD or DxxxD in most CDF proteins, such as in the Zn^2+^-transporting AtMTP1 and CrMTP1 (HxxxD), and in the Mn^2+^-transporting AtMTP11 and CrMTP4 (DxxxD), but the Group 5 TMD2 motif was A/SxxxD. Mutation of the His residue within this motif to Ala in the AtMTP1 Zn-CDF protein, the PtdMTP1 Zn-CDF protein from *Populus trichocarpa × deltoides* and the RmCzcD Zn-CDF protein from *Ralstonia metallidurans* all caused loss of function (Blaudez et al. 2003; Kawachi et al. 2012; Anton et al. 2004), perhaps indicating that these Group 5 proteins would not be functional for Zn^2+^ transport. However, there are exceptions to these motifs where His or Asp in the first position is not essential, such as NxxxD within TMD2 of the Mn^2+^ transporting SpMntA from *S. pneumoniae* or FxxxD within TMD2 of the Zn^2+^ transporting ScZRG17 from *S. cerevisiae* (Barber-Zucker et al. 2017). Furthermore, the recently identified Na^+^-transporting MceT protein from the halophilic bacterium *Planococcus dechangensis*, which has been proposed to be a member of a novel Na-CDF clade, has a distinct YxxxS motif (Xu et al. 2019). Currently we are unable to predict the substrate specificity of these Group 5 algae proteins from sequence comparisons therefore future experimental analysis is needed.

### C. reinhardtii MTP transporters have Zn-CDF and Mn-CDF functions

This study has identified *C. reinhardtii* members of the CDF family of metal efflux transporters as Zn^2+^, Co^2+^ and Mn^2+^ transporters, and validated the function of an algal Zn-CDF protein for the first time. CrMTP1 clearly groups in the Group 1 (Zn-CDF) clade while CrMTP2, CrMTP3 and CrMTP4 are clearly within the Group 4 (Mn-CDF) clade, and the yeast expression experiments validated these predicted activities, as we previously observed for CrMTP4 (Ibuot et al. 2017). An intriguing exception was the apparent Mn^2+^ and Zn^2+^ transport activity by CrMTP3, the first example we are aware of for a Mn^2+^-transporting CDF also being linked to Zn^2+^ transport. A Mn-CDF from *A. thaliana* (AtMTP8) may transport both Mn^2+^ and Fe^2+^ (Chu et al. 2017), while the Group 1 (Zn-CDF) member HsZnt10 can transport Mn^2+^ (Nishito et al. 2016). While there is some sequence variation between CrMTP3 and CrMTP2/CrMTP4 that could explain this difference, future biochemical and mutagenesis studies are required to experimental validate this further.

CrMTP1 was able to efficiently provide Zn tolerance to yeast, indicating Zn^2+^ transport activity, despite the structure of CrMTP1 being distinct from previously characterised Zn-CDF proteins such as AtMTP1 and ScZRC1 due to the significantly longer His-rich loop domain between TMD4 and 5. This seems to be a structural feature for many of the algal Zn-CDF proteins. The His-rich loop has been proposed to play a role in metal ion specificity and affinity, such that shortening of the AtMTP1 loop, which also reduces the number of His residues, alters the kinetics of Zn^2+^ transport, while the shortened loop or specific point-mutations within the loop can allow Co^2+^ transport, which is otherwise poor in native AtMTP1 (Kawachi et al. 2008; Podar et al. 2012). Therefore we might predict that the longer His-loop of CrMTP1 may provide different Zn^2+^ and Co^2+^ transport kinetics compared to some of the higher plant isoforms.

*CrMTP2* and *CrMTP4* but not the other *C. reinhardtii* CDF family genes have been shown to be transcriptionally induced by Mn deficiency, thus linking them further to Mn homeostasis roles (Allen et al. 2007), while *CrMTP1* is induced by Zn deficiency (Malasarn et al. 2013), although we could find no evidence by RT-PCR that any of these genes are transcriptionally regulated by metal excess. CrMTP1 was able to efficiently rescue the Zn sensitive phenotype of the yeast vacuolar Zn^2+^/H^+^ CDF transporter ScZRC1 and provide Zn tolerance in wild type yeast, thus it is proposed to provide a pathway of vacuolar Zn^2+^ sequestration equivalent to AtMTP1 and ScZRC1 (Kobae et al. 2004; MacDiarmid et al. 2002). Likewise, the significant Mn tolerance gained by CrMTP2, CrMTP3 or CrMTP4 that was coincident with increased Mn content within the cell is consistent with sequestration of Mn^2+^ into an internal compartment, possibly a vacuolar structure. *C. reinhardtii* lacks a large central vacuole as in typical plant cells but possess multiple lysosomal/vacuolar-like structures that are also referred to as acidocalcisomes (Blaby-Haas and Merchant 2014; Komine et al. 2000). These are acidic structures that accumulate polyphosphate and metals including Ca, Mg and Zn, and possess proton pumps that would be able to energise H^+^-coupled Zn^2+^ or Mn^2+^ transport by an MTP protein (Ruiz et al. 2001). The identification of the red algal MTP1 orthologue from *C. merolae* (Cmer_CMF058C) from an acidocalcisome proteomic study (Yagisawa et al. 2009), further strengthens the likely function of algal Zn-CDF proteins in vacuolar Zn^2+^ sequestration.

## Conclusions

Knowledge of sub-cellular metal transport processes, such as by members of the CDF transporter family, not only provides improved understanding of fundamental mechanisms of metal homeostasis and metal tolerance within a cell for a given organism, but may provide tools for biotechnological approaches to manipulate metal content within a whole organism, tissue, or sub-cellular compartment. Previous studies have proposed applications of CDF transporters for enhancing mineral biofortification of plants or for providing toxic metal bioremediation in plants or algae (Ibuot et al. 2017; Ricachenevsky et al. 2013; Das et al. 2016). This present study has indicated the wide diversity of CDF transporters that exist within the algae lineage. Although the vast majority of these proteins are still uncharacterized and the potential metal substrates are unknown, it is quite possible that some of these proteins will have characteristics that are distinct to those CDF transporters already known in plants and other eukaryotes. Therefore algal CDF genes may be attractive gene targets for future genetic engineering approaches in plants or microorganisms for metal transport engineering.

## Acknowledgements

This work was supported by a Government of Nigeria TETFUND PhD studentship awarded to AI. We thank Paul Lythgoe (Manchester Analytical Geochemical Unit, Department of Earth and Environmental Sciences, University of Manchester) for ICP-AES analysis. We thank Ute Krämer for providing the *zrc1 cot1* yeast strain.

