## Supplemental information for "Multi-genomic analysis of the Cation Diffusion Facilitator transporters from algae"

Aniefon Ibuot [a], Andrew P. Dean [b], and Jon K. Pittman [c,*]
[a] Department of Science Technology, Akwa Ibom State Polytechnic, Nigeria; [b] Department of Natural Sciences, Manchester Metropolitan University, UK; [c] Department of Earth and Environmental Sciences, The University of Manchester, UK

CDF protein sequence information was obtained from individual sequence database depositions followed published characterisation of selected CDF proteins or from genome annotations. Individual sequence, reference and database (including website address) information is provided in Supplementary Table S1 (below). All individual bacterial proteins sequences were obtained from National Center for Biotechnology Information (NCBI) Protein Database.

Genome assemblies and annotations from 38 species were used in this study. The details of each genome are described below. The majority of the genome data was accessed from the DOE Joint Genome Institute (JGI) Genome Portal (Grigoriev et al. 2012; Nordberg et al. 2014), including JGI MycoCosm (Grigoriev et al. 2014) and JGI PhycoCosm, and JGI Phyozome v.12 (Goodstein et al. 2012).

For the non-algal organisms the following genomes were analysed. *Nostoc punctiforme* PCC 73102 genome data was obtained from JGI Genome Portal. *Homo sapiens* sequence data was obtained from NCBI. *Saccharomyces cerevisiae* strain S288C genome data (genome version R64-2-1) was obtained from the *Saccharomyces* Genome Database (SGD) (Cherry et al. 2012; Engel et al. 2014). *Arabidopsis thaliana* cv. Col-0 genome (The Arabidopsis Genome Initiative 2000) release v.9 data was obtained from The Arabidopsis Information Resource (TAIR) v.10 (Lamesch et al. 2012). *Schizochytrium aggregatum* ATCC 28209 draft genome release v.1 data was obtained from JGI MycoCosm.

For a charophyte genome, *Klebsormidium nitens* NIES-2285 genome release v.1.1 data was obtained from the *Klebsormidium nitens* NIES-2285 genome project (Hori et al. 2014).

For the chlorophyte organisms the following genomes were analysed. *Bathycoccus prasinos* RCC1105 genome (Moreau et al. 2012) assembly ASM222023v1 data was obtained from JGI PhycoCosm. *Botryococcus braunii* Race B (Showa strain) genome release v.2.1 (Botryococcus braunii Showa v2.1 DOE-JGI, http://phytozome.jgi.doe.gov/) data was obtained from JGI Phytozome. *Chlamydomonas reinhardtii* CC-503 genome (Merchant et al. 2007) assembly and annotation v.5.5 data was obtained from JGI Phytozome. *Chlorella* sp. NC64A draft genome (Blanc et al. 2010) assembly and annotation v.1 data was obtained from JGI PhycoCosm. *Chromochloris zofingiensis* SAG 211-14 genome (Roth et al. 2017) assembly and annotation v.5.2.3.2 data was obtained from JGI PhycoCosm.*Coccomyxa subellipsoidea* C-169 genome (Blanc et al. 2012) assembly and annotation v.2 data was obtained from JGI Phytozome. *Dunaliella salina* CCAP19-18 draft genome (Polle et al. 2017) assembly and annotation v.1 data was obtained from JGI Phytozome. *Micromonas pusilla* CCMP1545 genome (Worden et al. 2009) assembly and annotation v.3 data was obtained from JGI Phytozome. *Micromonas* sp. RCC299 genome (Worden et al. 2009) assembly and annotation v.3 data was obtained from JGI Phytozome. *Ostreococcus lucimarinus* genome

(Palenik et al. 2007) assembly and annotation v.2 data was obtained from JGI Phytozome. *Ostreococcus* sp. RCC809 draft genome (Palenik et al. 2007) assembly and annotation v.2 data was obtained from JGI PhycoCosm.*Ostreococcus tauri* RCC4221 genome (Palenik et al. 2007; Blanc-Mathieu et al. 2014) assembly and annotation v.3 data was obtained from JGI PhycoCosm.*Volvox carteri* genome (Prochnik et al. 2010) assembly and annotation v.2.1 data was obtained from JGI Phytozome.

For the pelagophyte organisms the following genomes were analysed. *Aureococcus anophagefferens* clone 1984 genome (Gobler et al. 2011) assembly and annotation v.1 data was obtained from JGI PhycoCosm. *Pelagophyceae* sp. CCMP2097 draft genome assembly and annotation v.1 data was obtained from JGI PhycoCosm.

For the cryptophyte organisms the following genomes were analysed. *Cryptophyceae* sp. CCMP2293 draft genome assembly and annotation v.1 data was obtained from JGI PhycoCosm. *Guillardia theta* CCMP2712 draft genome (Curtis et al. 2012) assembly and annotation v.1 data was obtained from JGI PhycoCosm.

For the rhodophyte organisms the following genomes were analysed. *Cyanidioschyzon merolae* genome (Matsuzaki et al. 2004; Nozaki et al. 2007) assembly and annotation data was obtained from JGI PhycoCosm. *Galdieria sulphuraria* 074W genome (Schönknecht et al. 2013) assembly and annotation data was obtained from JGI PhycoCosm. *Porphyra umbilicalis* genome (Brawley et al. 2017) assembly and annotation v1.5 data was obtained from JGI Phytozome.

For the haptophyte organisms the following genomes were analysed. *Emiliania huxleyi* CCMP1516 draft genome (Read et al. 2013) assembly and annotation v.1 data was obtained from JGI PhycoCosm. *Pavlovales* sp. CCMP2436 draft genome assembly and annotation v.1 data was obtained from JGI PhycoCosm. *Phaeocystis antarctica* CCMP1374 genome assembly and annotation v.2.2 data was obtained from JGI PhycoCosm. *Phaeocystis globosa* Pg-G genome assembly and annotation v.2.3 data was obtained from JGI PhycoCosm.

For the bacillariophyte organisms the following genomes were analysed. *Fragilariopsis cylindrus* CCMP 1102 genome (Mock et al. 2017) assembly and annotation v.1 data was obtained from JGI PhycoCosm. *Phaeodactylum tricornutum* CCAP 1055/1 genome (Bowler et al. 2008) assembly and annotation v.2 data was obtained from JGI PhycoCosm. *Pseudo-nitzschia* multiseries CLN-47 draft genome assembly and annotation v.1 data was obtained from JGI PhycoCosm. *Thalassiosira pseudonana* CCMP 1335 genome (Armbrust et al. 2004) assembly and annotation v.3 data was obtained from JGI PhycoCosm.

For the ochrophyte organisms the following genomes were analysed. *Ectocarpus siliculosus* Ec 32 genome (Cock et al. 2010) assembly and annotation data was obtained from JGI PhycoCosm. *Nannochloropsis oceanica* CCMP1779 draft genome assembly and annotation v.2 data was obtained from JGI PhycoCosm. *Ochromonadaceae* sp. CCMP2298 draft genome assembly and annotation v.1 data was obtained from JGI PhycoCosm.

For a chlorarachinophyte genome, *Bigelowiella natans* CCMP2755 draft genome assembly and annotation v.1 data was obtained from JGI PhycoCosm.

**Supplementary Table S1.** Details of each CDF protein from each analysed organism and genome used in the phylogenetic analysis. The genome Id number or accession number refer to the database listed in the adjacent column, Database website addresses are shown in the table footnotes. CDF group assignments were determined from phylogenetic analysis (Figure 1) and supported by cluster analysis (Figure 2). Eight sequences were classed as unassigned including seven sequences where the evidence for Group 2 assignment was not supported by cluster analysis.

| Protein name | Species | Taxonomic classification | Genome ID or accession no. | Database | CDF Group | Known substrate | Reference |
|---|---|---|---|---|---|---|---|
| Bs_CzcD | *Bacillus subtilis* | Bacteria | NP_390542.1 | NCBI [a] | Group 1 (Zn-CDF) | $Zn^{2+}$, $Co^{2+}$, $Cu^{2+}$, $Ni^{2+}$ | Moore et al. (2005) |
| Bd_blr4938 | *Bradyrhizobium diazoefficiens* USDA 110 | Bacteria | NP_771578.1 | NCBI [a] | Group 1 (Zn-CDF) | | |
| Ec_ZitB | *Escherichia coli* | Bacteria | P75757.1 | NCBI [a] | Group 1 (Zn-CDF) | $Zn^{2+}$ | Chao & Fu (2004) |
| Ec_YiiP | *Escherichia coli* | Bacteria | P69380.1 | NCBI [a] | Group 2 (Fe/Zn CDF) | $Fe^{2+}$, $Zn^{2+}$, $Cd^{2+}$ | Wei & Fu (2005) |
| Cm_FieF | *Ralstonia metallidurans* CH34 | Bacteria | ZP_00593836.1 | NCBI [a] | Group 2 (Fe/Zn CDF) | $Fe^{2+}$, $Co^{2+}$, $Ni^{2+}$, $Cd^{2+}$, $Zn^{2+}$ | Munkelt et al. (2004) |
| Re_CepA | *Rhizobium etli* CFN 42 | Bacteria | ABC90024.1 | NCBI [a] | Group 1 (Zn-CDF) | $Co^{2+}$ | Cubillas et al. (2013) |
| Re_NepA | *Rhizobium etli* CFN 42 | Bacteria | ABC93653.1 | NCBI [a] | Group 2 (Fe/Zn CDF) | $Ni^{2+}$, $Co^{2+}$ | Cubillas et al. (2013) |
| Sm_0683 | *Sinorhizobium meliloti* 1021 | Bacteria | NP_435608.2 | NCBI [a] | Unassigned | | |
| Sp_MntE | *Streptococcus pneumoniae* D39 | Bacteria | ABJ55467.1 | NCBI [a] | Group 2 (Fe/Zn CDF) | $Mn^{2+}$ | Rosch et al. (2009) |
| Tt_CzrB | *Thermus thermophiles* | Bacteria | CAC83722.1 | NCBI [a] | Group 2 (Fe/Zn CDF) | $Zn^{2+}$, $Cd^{2+}$ | Spada et al. (2002) |
| Ma_4394 | *Methanosarcina acetivorans* C2A | Archaea | AAM07736.1 | NCBI [a] | Group 2 (Fe/Zn CDF) | | |
| Np_F1794 | *Nostoc punctiforme* PCC 73102 | Cyanobacteria | ACC80456.1 | JGI Genome Portal [b] | Group 3 (undefined) | | |
| Np_F0707 | *Nostoc punctiforme* PCC 73102 | Cyanobacteria | ACC79459.1 | JGI Genome Portal [b] | Group 2 (Fe/Zn CDF) | | |
| Np_R3104 | *Nostoc punctiforme* PCC 73102 | Cyanobacteria | ACC81585.1 | JGI Genome Portal [b] | Group 3 (undefined) | | |
| Np_F2455 | *Nostoc punctiforme* PCC 73102 | Cyanobacteria | ACC81016.1 | JGI Genome Portal [b] | Group 2 (Fe/Zn CDF) | | |
| Hs_Znt1 | *Homo sapiens* | Metazoa | NP_067017.2 | NCBI [a] | Group 1 (Zn-CDF) | $Zn^{2+}$ | Segal et al. (2004) |

| | | | | | | | |
|---|---|---|---|---|---|---|---|
| Hs_Znt2 | *Homo sapiens* | Metazoa | NP_001004434.1 | NCBI [a] | Group 1 (Zn-CDF) | $Zn^{2+}$ | Lopez & Kelleher (2009) |
| Hs_Znt3 | *Homo sapiens* | Metazoa | NP_003450.2 | NCBI [a] | Group 1 (Zn-CDF) | $Zn^{2+}$ | Salazar et al. (2009) |
| Hs_Znt4 | *Homo sapiens* | Metazoa | NP_037441.2 | NCBI [a] | Group 1 (Zn-CDF) | $Zn^{2+}$ | Murgia et al. (2011) |
| Hs_Znt5 | *Homo sapiens* | Metazoa | NP_075053.2 | NCBI [a] | Group 1 (Zn-CDF) | $Zn^{2+}$ | Ohana et al. (2009) |
| Hs_Znt6 | *Homo sapiens* | Metazoa | NP_001180442.1 | NCBI [a] | Group 1 (Zn-CDF) | $Zn^{2+}$ | Huang et al. (2002) |
| Hs_Znt7 | *Homo sapiens* | Metazoa | NP_001138356.1 | NCBI [a] | Group 1 (Zn-CDF) | $Zn^{2+}$ | Kirschke & Huang (2003) |
| Hs_Znt8 | *Homo sapiens* | Metazoa | NP_776250.2 | NCBI [a] | Group 1 (Zn-CDF) | $Zn^{2+}$ | Chimienti et al. (2004) |
| Hs_Znt9 | *Homo sapiens* | Metazoa | NP_006336.3 | NCBI [a] | Group 3 (undefined) | $Zn^{2+}$ | Perez et al. (2017) |
| Hs_Znt10 | *Homo sapiens* | Metazoa | NP_061183.2 | NCBI [a] | Group 1 (Zn-CDF) | $Mn^{2+}$, $Zn^{2+}$? | Nishito et al. (2016) |
| Sc_ZRC1 | *Saccharomyces cerevisiae* S288C | Ascomycota | SGD:S000004856 | SGC [c] | Group 1 (Zn-CDF) | $Zn^{2+}$ | MacDiarmid et al. (2002) |
| Sc_COT1 | *Saccharomyces cerevisiae* S288C | Ascomycota | SGD:S000005843 | SGC [c] | Group 1 (Zn-CDF) | $Zn^{2+}$, $Co^{2+}$ | MacDiarmid et al. (2002) |
| Sc_MSC2 | *Saccharomyces cerevisiae* S288C | Ascomycota | SGD:S000002613 | SGC [c] | Group 1 (Zn-CDF) | $Zn^{2+}$ | Ellis et al. (2004) |
| Sc_ZRG17 | *Saccharomyces cerevisiae* S288C | Ascomycota | SGD:S000005322 | SGC [c] | Unassigned | $Zn^{2+}$ | Ellis et al. (2005) |
| Sc_MMT1 | *Saccharomyces cerevisiae* S288C | Ascomycota | SGD:S000004789 | SGC [c] | Group 2 (Fe/Zn CDF) | $Fe^{2+}$ | Li et al. (2014) |
| Sc_MMT2 | *Saccharomyces cerevisiae* S288C | Ascomycota | SGD:S000006145 | SGC [c] | Group 2 (Fe/Zn CDF) | $Fe^{2+}$ | Li et al. (2014) |
| At_MTP1 | *Arabidopsis thaliana* columbia | Embryophyta | AT2G46800.1 | TAIR [d] | Group 1 (Zn-CDF) | $Zn^{2+}$ | Kobae et al. (2004) |
| At_MTP2 | *Arabidopsis thaliana* columbia | Embryophyta | AT3G61940.1 | TAIR [d] | Group 1 (Zn-CDF) | $Zn^{2+}$ | Sinclair et al. (2018) |
| At_MTP3 | *Arabidopsis thaliana* columbia | Embryophyta | AT3G58810.1 | TAIR [d] | Group 1 (Zn-CDF) | $Zn^{2+}$, $Cd^{2+}$ | Arrivault et al. (2006) |
| At_MTP4 | *Arabidopsis thaliana* columbia | Embryophyta | AT2G29410.1 | TAIR [d] | Group 1 (Zn-CDF) | $Zn^{2+}$ | Fujiwara et al. (2015) |
| At_MTP5 | *Arabidopsis thaliana* columbia | Embryophyta | AT3G12100.1 | TAIR [d] | Group 1 (Zn-CDF) | $Zn^{2+}$ | Fujiwara et al. (2015) |

| | | | | | | | |
|---|---|---|---|---|---|---|---|
| At_MTP6 | *Arabidopsis thaliana* columbia | Embryophyta | AT2G47830.1 | TAIR [d] | Group 2 (Fe/Zn CDF) | | |
| At_MTP7 | *Arabidopsis thaliana* columbia | Embryophyta | AT1G51610.1 | TAIR [d] | Group 3 (undefined) | | |
| At_MTP8 | *Arabidopsis thaliana* columbia | Embryophyta | AT3G58060.1 | TAIR [d] | Group 4 (Mn-CDF) | $Mn^{2+}$, $Fe^{2+}$? | Eroglu et al. (2016) |
| At_MTP9 | *Arabidopsis thaliana* columbia | Embryophyta | AT1G79520.2 | TAIR [d] | Group 4 (Mn-CDF) | $Mn^{2+}$ | Chu et al. (2017) |
| At_MTP10 | *Arabidopsis thaliana* columbia | Embryophyta | AT1G16310.1 | TAIR [d] | Group 4 (Mn-CDF) | $Mn^{2+}$ | Chu et al. (2017) |
| At_MTP11 | *Arabidopsis thaliana* columbia | Embryophyta | AT2G39450.1 | TAIR [d] | Group 4 (Mn-CDF) | $Mn^{2+}$ | Peiter et al. (2007) |
| At_MTP12 | *Arabidopsis thaliana* columbia | Embryophyta | AT2G04620.1 | TAIR [d] | Group 1 (Zn-CDF) | $Zn^{2+}$ | Fujiwara et al. (2015) |
| Sa_54404 | *Schizochytrium aggregatum* ATCC 28209 | Thraustochytrid | Protein Id: 54404 | JGI MycoCosm [e] | Group 1 (Zn-CDF) | | |
| Sa_80149 | *Schizochytrium aggregatum* ATCC 28209 | Thraustochytrid | Protein Id: 80149 | JGI MycoCosm [e] | Group 1 (Zn-CDF) | | |
| Sa_81417 | *Schizochytrium aggregatum* ATCC 28209 | Thraustochytrid | Protein Id: 81417 | JGI MycoCosm [e] | Group 1 (Zn-CDF) | | |
| Sa_27533 | *Schizochytrium aggregatum* ATCC 28209 | Thraustochytrid | Protein Id: 27533 | JGI MycoCosm [e] | Group 2 (Fe/Zn CDF) | | |
| Sa_73731 | *Schizochytrium aggregatum* ATCC 28209 | Thraustochytrid | Protein Id: 73731 | JGI MycoCosm [e] | Group 3 (undefined) | | |
| Sa_13553 | *Schizochytrium aggregatum* ATCC 28209 | Thraustochytrid | Protein Id: 13553 | JGI MycoCosm [e] | Group 4 (Mn-CDF) | | |
| Sa_86437 | *Schizochytrium aggregatum* ATCC 28209 | Thraustochytrid | Protein Id: 86437 | JGI MycoCosm [e] | Group 5 (undefined) | | |
| Kn_00085_0150 | *Klebsormidium nitens* NIES-2285 | Charophyta | kfl00085_0150_v1.1 | Klebsormidium genome project [f] | Group 1 (Zn-CDF) | | |
| Kn_00345_0040 | *Klebsormidium nitens* NIES-2285 | Charophyta | kfl00345_0040_v1.1 | Klebsormidium genome project [f] | Group 1 (Zn-CDF) | | |
| Kn_00701_0020 | *Klebsormidium nitens* NIES-2285 | Charophyta | kfl00701_0020_v1.1 | Klebsormidium genome project [f] | Group 1 (Zn-CDF) | | |
| Kn_00804_0020 | *Klebsormidium nitens* NIES-2285 | Charophyta | kfl00804_0020_v1.1 | Klebsormidium genome project [f] | Group 1 (Zn-CDF) | | |
| Kn_00270_0190 | *Klebsormidium nitens* NIES-2285 | Charophyta | kfl00270_0190_v1.1 | Klebsormidium genome project [f] | Group 2 (Fe/Zn CDF) | | |
| Kn_00187_0105 | *Klebsormidium nitens* NIES-2285 | Charophyta | kfl00187_0105_v1.1 | Klebsormidium genome project [f] | Group 3 (undefined) | | |

| | | | | | | | |
|---|---|---|---|---|---|---|---|
| Kn_00078_0240 | *Klebsormidium nitens* NIES-2285 | Charophyta | kfl00078_0240_v1.1 | Klebsormidium genome project [f] | Group 4 (Mn-CDF) | | |
| Kn_00343_0090 | *Klebsormidium nitens* NIES-2285 | Charophyta | kfl00343_0090_v1.1 | Klebsormidium genome project [f] | Group 4 (Mn-CDF) | | |
| Bp_6660 | *Bathycoccus prasinos* RCC1105 | Chlorophyta | Protein Id: 6660 | JGI PhycoCosm [g] | Group 2 (Fe/Zn CDF) | | |
| Bp_127 | *Bathycoccus prasinos* RCC1105 | Chlorophyta | Protein Id: 127 | JGI PhycoCosm [g] | Group 3 (undefined) | | |
| Bp_7715 | *Bathycoccus prasinos* RCC1105 | Chlorophyta | Protein Id: 7715 | JGI PhycoCosm [g] | Group 4 (Mn-CDF) | | |
| Bb_0249s0060 | *Botryococcus braunii* Race B (Showa strain) | Chlorophyta | Bobra.0249s0060.1 | JGI Phytozome [h] | Group 1 (Zn-CDF) | | |
| Bb_0056s0042 | *Botryococcus braunii* Race B (Showa strain) | Chlorophyta | Bobra.0056s0042.1 | JGI Phytozome [h] | Group 3 (undefined) | | |
| Cr_MTP1 | *Chlamydomonas reinhardtii* CC-503 | Chlorophyta | Cre03.g145087.t1.1 | JGI Phytozome [h] | Group 1 (Zn-CDF) | $Zn^{2+}$, $Co^{2+}$ | This study |
| Cr_MTP2 | *Chlamydomonas reinhardtii* CC-503 | Chlorophyta | Cre03.g160800.t1.2 | JGI Phytozome [h] | Group 4 (Mn-CDF) | $Mn^{2+}$ | This study |
| Cr_MTP3 | *Chlamydomonas reinhardtii* CC-503 | Chlorophyta | Cre03.g160750.t1.1 | JGI Phytozome [h] | Group 4 (Mn-CDF) | $Mn^{2+}$, $Zn^{2+}$, $Co^{2+}$ | This study |
| Cr_MTP4 | *Chlamydomonas reinhardtii* CC-503 | Chlorophyta | Cre03.g160550.t1.1 | JGI Phytozome [h] | Group 4 (Mn-CDF) | $Mn^{2+}$, $Cd^{2+}$ | Ibuot et al. (2017) |
| Cr_MTP5 | *Chlamydomonas reinhardtii* CC-503 | Chlorophyta | Cre06.g289150.t1.1 | JGI Phytozome [h] | Group 3 (undefined) | | |
| Chlsp_37115 | *Chlorella* sp. NC64A | Chlorophyta | Protein Id: 37115 | JGI PhycoCosm [g] | Group 1 (Zn-CDF) | | |
| Chlsp_18700 | *Chlorella* sp. NC64A | Chlorophyta | Protein Id: 18700 | JGI PhycoCosm [g] | Group 3 (undefined) | | |
| Chlsp_59655 | *Chlorella* sp. NC64A | Chlorophyta | Protein Id: 59655 | JGI PhycoCosm [g] | Group 4 (Mn-CDF) | | |
| Cz_04g25040 | *Chromochloris zofingiensis* SAG 211-14 | Chlorophyta | Cz04g25040.t1 | JGI PhycoCosm [g] | Group 1 (Zn-CDF) | | |
| Cz_11g13120 | *Chromochloris zofingiensis* SAG 211-14 | Chlorophyta | Cz11g13120.t1 | JGI PhycoCosm [g] | Group 3 (undefined) | | |
| Cz_12g10240 | *Chromochloris zofingiensis* SAG 211-14 | Chlorophyta | Cz12g10240.t1 | JGI PhycoCosm [g] | Group 4 (Mn-CDF) | | |
| Cz_Ug00396 | *Chromochloris zofingiensis* SAG 211-14 | Chlorophyta | UNPLg00396.t1 | JGI PhycoCosm [g] | Group 4 (Mn-CDF) | | |
| Cs_14234 | *Coccomyxa subellipsoidea* C-169 | Chlorophyta | Protein Id: 14234 | JGI Phytozome [h] | Group 1 (Zn-CDF) | | |

| | | | | | |
|---|---|---|---|---|---|
| Cs_16095 | *Coccomyxa subellipsoidea* C-169 | Chlorophyta | Protein Id: 16095 | JGI Phytozome [h] | Group 1 (Zn-CDF) |
| Cs_28593 | *Coccomyxa subellipsoidea* C-169 | Chlorophyta | Protein Id: 28593 | JGI Phytozome [h] | Group 3 (undefined) |
| Cs_21325 | *Coccomyxa subellipsoidea* C-169 | Chlorophyta | Protein Id: 21325 | JGI Phytozome [h] | Group 4 (Mn-CDF) |
| Cs_39175 | *Coccomyxa subellipsoidea* C-169 | Chlorophyta | Protein Id: 39175 | JGI Phytozome [h] | Group 4 (Mn-CDF) |
| Ds_0333s00001 | *Dunaliella salina* CCAP19-18 | Chlorophyta | Dusal.0333s00001.1 | JGI Phytozome [h] | Group 1 (Zn-CDF) |
| Ds_0333s00003 | *Dunaliella salina* CCAP19-18 | Chlorophyta | Dusal.0333s00003.1 | JGI Phytozome [h] | Group 1 (Zn-CDF) |
| Ds_1360s00001 | *Dunaliella salina* CCAP19-18 | Chlorophyta | Dusal.1360s00001.1 | JGI Phytozome [h] | Group 3 (undefined) |
| Ds_0199s00009 | *Dunaliella salina* CCAP19-18 | Chlorophyta | Dusal.0199s00009.1 | JGI Phytozome [h] | Group 4 (Mn-CDF) |
| Mp_17485 | *Micromonas pusilla* CCMP1545 | Chlorophyta | Protein Id: 17485 | JGI Phytozome [h] | Group 3 (undefined) |
| Mp_52662 | *Micromonas pusilla* CCMP1545 | Chlorophyta | Protein Id: 52662 | JGI Phytozome [h] | Group 4 (Mn-CDF) |
| Msp_60348 | *Micromonas* sp. RCC299 | Chlorophyta | Protein Id: 60348 | JGI Phytozome [h] | Group 1 (Zn-CDF) |
| Msp_102742 | *Micromonas* sp. RCC299 | Chlorophyta | Protein Id: 102742 | JGI Phytozome [h] | Group 1 (Zn-CDF) |
| Msp_64402 | *Micromonas* sp. RCC299 | Chlorophyta | Protein Id: 64402 | JGI Phytozome [h] | Group 2 (Fe/Zn CDF) |
| Msp_97795 | *Micromonas* sp. RCC299 | Chlorophyta | Protein Id: 97795 | JGI Phytozome [h] | Group 3 (undefined) |
| Msp_55479 | *Micromonas* sp. RCC299 | Chlorophyta | Protein Id: 55479 | JGI Phytozome [h] | Group 4 (Mn-CDF) |
| Ol_39741 | *Ostreococcus lucimarinus* | Chlorophyta | Protein Id: 39741 | JGI Phytozome [h] | Group 3 (undefined) |
| Ol_33926 | *Ostreococcus lucimarinus* | Chlorophyta | Protein Id: 33926 | JGI Phytozome [h] | Group 4 (Mn-CDF) |
| Ostsp_28125 | *Ostreococcus* sp. RCC809 | Chlorophyta | Protein Id: 28125 | JGI PhycoCosm [g] | Group 3 (undefined) |
| Ostsp_30303 | *Ostreococcus* sp. RCC809 | Chlorophyta | Protein Id: 30303 | JGI PhycoCosm [g] | Group 4 (Mn-CDF) |
| Ot_69157 | *Ostreococcus tauri* RCC4221 | Chlorophyta | Protein Id: 69157 | JGI PhycoCosm [g] | Group 3 (undefined) |

| | | | | | |
|---|---|---|---|---|---|
| Ot_73435 | *Ostreococcus tauri* RCC4221 | Chlorophyta | Protein Id: 73435 | JGI PhycoCosm [g] | Group 4 (Mn-CDF) |
| Vc_0014s0072 | *Volvox carteri* | Chlorophyta | Vocar.0014s0072.1 | JGI Phytozome [h] | Group 1 (Zn-CDF) |
| Vc_0063s0011 | *Volvox carteri* | Chlorophyta | Vocar.0063s0011.1 | JGI Phytozome [h] | Group 3 (undefined) |
| Vc_0005s0300 | *Volvox carteri* | Chlorophyta | Vocar.0005s0300.1 | JGI Phytozome [h] | Group 4 (Mn-CDF) |
| Aa_18887 | *Aureococcus anophagefferens* clone 1984 | Pelagophyta | Protein Id: 18887 | JGI PhycoCosm [g] | Group 1 (Zn-CDF) |
| Aa_54063 | *Aureococcus anophagefferens* clone 1984 | Pelagophyta | Protein Id: 54063 | JGI PhycoCosm [g] | Group 2 (Fe/Zn CDF) |
| Aa_63403 | *Aureococcus anophagefferens* clone 1984 | Pelagophyta | Protein Id: 63403 | JGI PhycoCosm [g] | Group 2 (Fe/Zn CDF) |
| Aa_12227 | *Aureococcus anophagefferens* clone 1984 | Pelagophyta | Protein Id: 12227 | JGI PhycoCosm [g] | Group 4 (Mn-CDF) |
| Psp_582096 | *Pelagophyceae* sp. CCMP2097 | Pelagophyta | Protein Id: 582096 | JGI PhycoCosm [g] | Group 1 (Zn-CDF) |
| Psp_474385 | *Pelagophyceae* sp. CCMP2097 | Pelagophyta | Protein Id: 474385 | JGI PhycoCosm [g] | Group 2 (Fe/Zn CDF) |
| Psp_505673 | *Pelagophyceae* sp. CCMP2097 | Pelagophyta | Protein Id: 505673 | JGI PhycoCosm [g] | Group 2 (Fe/Zn CDF) |
| Psp_623472 | *Pelagophyceae* sp. CCMP2097 | Pelagophyta | Protein Id: 623472 | JGI PhycoCosm [g] | Group 3 (undefined) |
| Psp_507764 | *Pelagophyceae* sp. CCMP2097 | Pelagophyta | Protein Id: 507764 | JGI PhycoCosm [g] | Group 4 (Mn-CDF) |
| Psp_522664 | *Pelagophyceae* sp. CCMP2097 | Pelagophyta | Protein Id: 522664 | JGI PhycoCosm [g] | Group 4 (Mn-CDF) |
| Psp_509680 | *Pelagophyceae* sp. CCMP2097 | Pelagophyta | Protein Id: 509680 | JGI PhycoCosm [g] | Group 5 (undefined) |
| Psp_614657 | *Pelagophyceae* sp. CCMP2097 | Pelagophyta | Protein Id: 614657 | JGI PhycoCosm [g] | Group 5 (undefined) |
| Psp_615359 | *Pelagophyceae* sp. CCMP2097 | Pelagophyta | Protein Id: 615359 | JGI PhycoCosm [g] | Group 5 (undefined) |
| Psp_617317 | *Pelagophyceae* sp. CCMP2097 | Pelagophyta | Protein Id: 617317 | JGI PhycoCosm [g] | Group 5 (undefined) |

| | | | | | |
|---|---|---|---|---|---|
| Psp_584138 | *Pelagophyceae* sp. CCMP2097 | Pelagophyta | Protein Id: 584138 | JGI PhycoCosm [g] | Unassigned |
| Csp_1897108 | *Cryptophyceae* sp. CCMP2293 | Cryptophyta | Protein Id: 1897108 | JGI PhycoCosm [g] | Group 1 (Zn-CDF) |
| Csp_1763656 | *Cryptophyceae* sp. CCMP2293 | Cryptophyta | Protein Id: 1763656 | JGI PhycoCosm [g] | Group 2 (Fe/Zn CDF) |
| Csp_1785488 | *Cryptophyceae* sp. CCMP2293 | Cryptophyta | Protein Id: 1785488 | JGI PhycoCosm [g] | Group 5 (undefined) |
| Gt_65516 | *Guillardia theta* CCMP2712 | Cryptophyta | Protein Id: 65516 | JGI PhycoCosm [g] | Group 1 (Zn-CDF) |
| Gt_131845 | *Guillardia theta* CCMP2712 | Cryptophyta | Protein Id: 131845 | JGI PhycoCosm [g] | Group 1 (Zn-CDF) |
| Gt_133916 | *Guillardia theta* CCMP2712 | Cryptophyta | Protein Id: 133916 | JGI PhycoCosm [g] | Group 1 (Zn-CDF) |
| Gt_140741 | *Guillardia theta* CCMP2712 | Cryptophyta | Protein Id: 140741 | JGI PhycoCosm [g] | Group 1 (Zn-CDF) |
| Gt_163374 | *Guillardia theta* CCMP2712 | Cryptophyta | Protein Id: 163374 | JGI PhycoCosm [g] | Group 1 (Zn-CDF) |
| Gt_163373 | *Guillardia theta* CCMP2712 | Cryptophyta | Protein Id: 163373 | JGI PhycoCosm [g] | Group 1 (Zn-CDF) |
| Gt_137697 | *Guillardia theta* CCMP2712 | Cryptophyta | Protein Id: 137697 | JGI PhycoCosm [g] | Group 2 (Fe/Zn CDF) |
| Gt_114253 | *Guillardia theta* CCMP2712 | Cryptophyta | Protein Id: 114253 | JGI PhycoCosm [g] | Group 3 (undefined) |
| Gt_77758 | *Guillardia theta* CCMP2712 | Cryptophyta | Protein Id: 77758 | JGI PhycoCosm [g] | Group 4 (Mn-CDF) |
| Gt_165502 | *Guillardia theta* CCMP2712 | Cryptophyta | Protein Id: 165502 | JGI PhycoCosm [g] | Group 5 (undefined) |
| Cmer_CMF058C | *Cyanidioschyzon merolae* strain 10D | Rhodophyta | CMF058C | JGI PhycoCosm [g] | Group 1 (Zn-CDF) |
| Cmer_CMT536C | *Cyanidioschyzon merolae* strain 10D | Rhodophyta | CMT536C | JGI PhycoCosm [g] | Group 1 (Zn-CDF) |
| Cmer_CMC075C | *Cyanidioschyzon merolae* strain 10D | Rhodophyta | CMC075C | JGI PhycoCosm [g] | Group 4 (Mn-CDF) |
| Gs_1521 | *Galdieria sulphuraria* 074W | Rhodophyta | Protein Id: 1521 | JGI PhycoCosm [g] | Group 1 (Zn-CDF) |
| Pu_0152s0012 | *Porphyra umbilicalis* | Rhodophyta | Pum0152s0012.1 | JGI Phytozome [h] | Group 1 (Zn-CDF) |
| Pu_0222s0004 | *Porphyra umbilicalis* | Rhodophyta | Pum0222s0004.1 | JGI Phytozome [h] | Group 1 (Zn-CDF) |

| | | | | | |
|---|---|---|---|---|---|
| Pu_0333s0033 | *Porphyra umbilicalis* | Rhodophyta | Pum0333s0033.1 | JGI Phytozome [h] | Group 1 (Zn-CDF) |
| Pu_0056s0045 | *Porphyra umbilicalis* | Rhodophyta | Pum0056s0045.1 | JGI Phytozome [h] | Group 4 (Mn-CDF) |
| Eh_42691 | *Emiliania huxleyi* CCMP1516 | Haptophyta | Protein Id: 42691 | JGI PhycoCosm [g] | Group 1 (Zn-CDF) |
| Eh_53810 | *Emiliania huxleyi* CCMP1516 | Haptophyta | Protein Id: 53810 | JGI PhycoCosm [g] | Group 1 (Zn-CDF) |
| Eh_414632 | *Emiliania huxleyi* CCMP1516 | Haptophyta | Protein Id: 414632 | JGI PhycoCosm [g] | Group 1 (Zn-CDF) |
| Eh_98399 | *Emiliania huxleyi* CCMP1516 | Haptophyta | Protein Id: 98399 | JGI PhycoCosm [g] | Group 2 (Fe/Zn CDF) |
| Eh_202761 | *Emiliania huxleyi* CCMP1516 | Haptophyta | Protein Id: 202761 | JGI PhycoCosm [g] | Group 2 (Fe/Zn CDF) |
| Eh_210717 | *Emiliania huxleyi* CCMP1516 | Haptophyta | Protein Id: 210717 | JGI PhycoCosm [g] | Group 2 (Fe/Zn CDF) |
| Eh_461752 | *Emiliania huxleyi* CCMP1516 | Haptophyta | Protein Id: 461752 | JGI PhycoCosm [g] | Group 2 (Fe/Zn CDF) |
| Eh_42214 | *Emiliania huxleyi* CCMP1516 | Haptophyta | Protein Id: 42214 | JGI PhycoCosm [g] | Group 4 (Mn-CDF) |
| Eh_117787 | *Emiliania huxleyi* CCMP1516 | Haptophyta | Protein Id: 117787 | JGI PhycoCosm [g] | Group 5 (undefined) |
| Eh_203258 | *Emiliania huxleyi* CCMP1516 | Haptophyta | Protein Id: 203258 | JGI PhycoCosm [g] | Group 5 (undefined) |
| Eh_211062 | *Emiliania huxleyi* CCMP1516 | Haptophyta | Protein Id: 211062 | JGI PhycoCosm [g] | Group 5 (undefined) |
| Pasp_262010 | *Pavlovales* sp. CCMP2436 | Haptophyta | Protein Id: 262010 | JGI PhycoCosm [g] | Group 1 (Zn-CDF) |
| Pasp_936591 | *Pavlovales* sp. CCMP2436 | Haptophyta | Protein Id: 936591 | JGI PhycoCosm [g] | Group 1 (Zn-CDF) |
| Pasp_580610 | *Pavlovales* sp. CCMP2436 | Haptophyta | Protein Id: 580610 | JGI PhycoCosm [g] | Group 2 (Fe/Zn CDF) |
| Pasp_597035 | *Pavlovales* sp. CCMP2436 | Haptophyta | Protein Id: 597035 | JGI PhycoCosm [g] | Group 2 (Fe/Zn CDF) |
| Pasp_407315 | *Pavlovales* sp. CCMP2436 | Haptophyta | Protein Id: 407315 | JGI PhycoCosm [g] | Group 4 (Mn-CDF) |
| Pasp_915848 | *Pavlovales* sp. CCMP2436 | Haptophyta | Protein Id: 915848 | JGI PhycoCosm [g] | Group 4 (Mn-CDF) |
| Pasp_939555 | *Pavlovales* sp. CCMP2436 | Haptophyta | Protein Id: 939555 | JGI PhycoCosm [g] | Group 5 (undefined) |

| | | | | | |
|---|---|---|---|---|---|
| Pasp_1059224 | *Pavlovales* sp. CCMP2436 | Haptophyta | Protein Id: 1059224 | JGI PhycoCosm [g] | Group 5 (undefined) |
| Pasp_1061345 | *Pavlovales* sp. CCMP2436 | Haptophyta | Protein Id: 1061345 | JGI PhycoCosm [g] | Group 5 (undefined) |
| Pa_1122 | *Phaeocystis antarctica* CCMP1374 | Haptophyta | Protein Id: 1122 | JGI PhycoCosm [g] | Group 1 (Zn-CDF) |
| Pa_9803 | *Phaeocystis antarctica* CCMP1374 | Haptophyta | Protein Id: 9803 | JGI PhycoCosm [g] | Group 1 (Zn-CDF) |
| Pa_16275 | *Phaeocystis antarctica* CCMP1374 | Haptophyta | Protein Id: 16275 | JGI PhycoCosm [g] | Group 1 (Zn-CDF) |
| Pa_24855 | *Phaeocystis antarctica* CCMP1374 | Haptophyta | Protein Id: 24855 | JGI PhycoCosm [g] | Group 1 (Zn-CDF) |
| Pa_30362 | *Phaeocystis antarctica* CCMP1374 | Haptophyta | Protein Id: 30362 | JGI PhycoCosm [g] | Group 1 (Zn-CDF) |
| Pa_1127 | *Phaeocystis antarctica* CCMP1374 | Haptophyta | Protein Id: 1127 | JGI PhycoCosm [g] | Group 2 (Fe/Zn CDF) |
| Pa_2008 | *Phaeocystis antarctica* CCMP1374 | Haptophyta | Protein Id: 2008 | JGI PhycoCosm [g] | Group 2 (Fe/Zn CDF) |
| Pa_9269 | *Phaeocystis antarctica* CCMP1374 | Haptophyta | Protein Id: 9269 | JGI PhycoCosm [g] | Group 2 (Fe/Zn CDF) |
| Pa_17347 | *Phaeocystis antarctica* CCMP1374 | Haptophyta | Protein Id: 17347 | JGI PhycoCosm [g] | Group 2 (Fe/Zn CDF) |
| Pa_17350 | *Phaeocystis antarctica* CCMP1374 | Haptophyta | Protein Id: 17350 | JGI PhycoCosm [g] | Group 2 (Fe/Zn CDF) |
| Pa_19534 | *Phaeocystis antarctica* CCMP1374 | Haptophyta | Protein Id: 19534 | JGI PhycoCosm [g] | Group 2 (Fe/Zn CDF) |
| Pa_27928 | *Phaeocystis antarctica* CCMP1374 | Haptophyta | Protein Id: 27928 | JGI PhycoCosm [g] | Group 2 (Fe/Zn CDF) |
| Pa_35928 | *Phaeocystis antarctica* CCMP1374 | Haptophyta | Protein Id: 35928 | JGI PhycoCosm [g] | Group 3 (undefined) |
| Pa_24952 | *Phaeocystis antarctica* CCMP1374 | Haptophyta | Protein Id: 24952 | JGI PhycoCosm [g] | Group 4 (Mn-CDF) |
| Pg_215 | *Phaeocystis globosa* Pg-G | Haptophyta | Protein Id: 215 | JGI PhycoCosm [g] | Group 1 (Zn-CDF) |
| Pg_11101 | *Phaeocystis globosa* Pg-G | Haptophyta | Protein Id: 11101 | JGI PhycoCosm [g] | Group 1 (Zn-CDF) |
| Pg_2185 | *Phaeocystis globosa* Pg-G | Haptophyta | Protein Id: 2185 | JGI PhycoCosm [g] | Group 2 (Fe/Zn CDF) |
| Pg_4331 | *Phaeocystis globosa* Pg-G | Haptophyta | Protein Id: 4331 | JGI PhycoCosm [g] | Group 2 (Fe/Zn CDF) |

| | | | | | |
|---|---|---|---|---|---|
| Pg_30462 | *Phaeocystis globosa* Pg-G | Haptophyta | Protein Id: 30462 | JGI PhycoCosm [g] | Group 2 (Fe/Zn CDF) |
| Pg_25361 | *Phaeocystis globosa* Pg-G | Haptophyta | Protein Id: 25361 | JGI PhycoCosm [g] | Group 2 (Fe/Zn CDF) |
| Pg_17382 | *Phaeocystis globosa* Pg-G | Haptophyta | Protein Id: 17382 | JGI PhycoCosm [g] | Group 4 (Mn-CDF) |
| Pg_2642 | *Phaeocystis globosa* Pg-G | Haptophyta | Protein Id: 2642 | JGI PhycoCosm [g] | Group 5 (undefined) |
| Pg_5451 | *Phaeocystis globosa* Pg-G | Haptophyta | Protein Id: 5451 | JGI PhycoCosm [g] | Group 5 (undefined) |
| Fc_185721 | *Fragilariopsis cylindrus* CCMP 1102 | Bacillariophyta | Protein Id: 185721 | JGI PhycoCosm [g] | Group 1 (Zn-CDF) |
| Fc_197703 | *Fragilariopsis cylindrus* CCMP 1102 | Bacillariophyta | Protein Id: 197703 | JGI PhycoCosm [g] | Group 2 (Fe/Zn CDF) |
| Fc_138643 | *Fragilariopsis cylindrus* CCMP 1102 | Bacillariophyta | Protein Id: 138643 | JGI PhycoCosm [g] | Group 3 (undefined) |
| Fc_179287 | *Fragilariopsis cylindrus* CCMP 1102 | Bacillariophyta | Protein Id: 179287 | JGI PhycoCosm [g] | Group 5 (undefined) |
| Fc_243722 | *Fragilariopsis cylindrus* CCMP 1102 | Bacillariophyta | Protein Id: 243722 | JGI PhycoCosm [g] | Group 5 (undefined) |
| Fc_244237 | *Fragilariopsis cylindrus* CCMP 1102 | Bacillariophyta | Protein Id: 244237 | JGI PhycoCosm [g] | Group 5 (undefined) |
| Pt_23557 | *Phaeodactylum tricornutum* CCAP 1055/1 | Bacillariophyta | Protein Id: 23557 | JGI PhycoCosm [g] | Group 1 (Zn-CDF) |
| Pt_45119 | *Phaeodactylum tricornutum* CCAP 1055/1 | Bacillariophyta | Protein Id: 45119 | JGI PhycoCosm [g] | Group 2 (Fe/Zn CDF) |
| Pt_13994 | *Phaeodactylum tricornutum* CCAP 1055/1 | Bacillariophyta | Protein Id: 13994 | JGI PhycoCosm [g] | Group 3 (undefined) |
| Pt_48955 | *Phaeodactylum tricornutum* CCAP 1055/1 | Bacillariophyta | Protein Id: 48955 | JGI PhycoCosm [g] | Group 4 (Mn-CDF) |
| Pt_46297 | *Phaeodactylum tricornutum* CCAP 1055/1 | Bacillariophyta | Protein Id: 46297 | JGI PhycoCosm [g] | Group 5 (undefined) |
| Pn_98747 | *Pseudo-nitzschia* multiseries CLN-47 | Bacillariophyta | Protein Id: 98747 | JGI PhycoCosm [g] | Group 1 (Zn-CDF) |
| Pn_260498 | *Pseudo-nitzschia* multiseries CLN-47 | Bacillariophyta | Protein Id: 260498 | JGI PhycoCosm [g] | Group 1 (Zn-CDF) |
| Pn_290477 | *Pseudo-nitzschia* multiseries CLN-47 | Bacillariophyta | Protein Id: 290477 | JGI PhycoCosm [g] | Group 2 (Fe/Zn CDF) |
| Pn_247016 | *Pseudo-nitzschia* multiseries CLN-47 | Bacillariophyta | Protein Id: 247016 | JGI PhycoCosm [g] | Group 3 (undefined) |

| | | | | | |
|---|---|---|---|---|---|
| Pn_153478 | *Pseudo-nitzschia* multiseries CLN-47 | Bacillariophyta | Protein Id: 153478 | JGI PhycoCosm [g] | Group 4 (Mn-CDF) |
| Pn_302540 | *Pseudo-nitzschia* multiseries CLN-47 | Bacillariophyta | Protein Id: 302540 | JGI PhycoCosm [g] | Group 5 (undefined) |
| Tp_26530 | *Thalassiosira pseudonana* CCMP 1335 | Bacillariophyta | Protein Id: 26530 | JGI PhycoCosm [g] | Group 1 (Zn-CDF) |
| Tp_260789 | *Thalassiosira pseudonana* CCMP 1335 | Bacillariophyta | Protein Id: 260789 | JGI PhycoCosm [g] | Group 2 (Fe/Zn CDF) |
| Tp_7978 | *Thalassiosira pseudonana* CCMP 1335 | Bacillariophyta | Protein Id: 7978 | JGI PhycoCosm [g] | Group 3 (undefined) |
| Tp_31951 | *Thalassiosira pseudonana* CCMP 1335 | Bacillariophyta | Protein Id: 31951 | JGI PhycoCosm [g] | Group 4 (Mn-CDF) |
| Tp_22171 | *Thalassiosira pseudonana* CCMP 1335 | Bacillariophyta | Protein Id: 22171 | JGI PhycoCosm [g] | Unassigned |
| Es_24050 | *Ectocarpus siliculosus* Ec 32 | Ochrophyta | Protein Id: 24050 | JGI PhycoCosm [g] | Group 1 (Zn-CDF) |
| Es_26403 | *Ectocarpus siliculosus* Ec 32 | Ochrophyta | Protein Id: 26403 | JGI PhycoCosm [g] | Group 2 (Fe/Zn CDF) |
| Es_21867 | *Ectocarpus siliculosus* Ec 32 | Ochrophyta | Protein Id: 21867 | JGI PhycoCosm [g] | Group 3 (undefined) |
| Es_29376 | *Ectocarpus siliculosus* Ec 32 | Ochrophyta | Protein Id: 29376 | JGI PhycoCosm [g] | Group 5 (undefined) |
| Es_18100 | *Ectocarpus siliculosus* Ec 32 | Ochrophyta | Protein Id: 18100 | JGI PhycoCosm [g] | Unassigned |
| No_630191 | *Nannochloropsis oceanica* CCMP1779 | Ochrophyta | Protein Id: 630191 | JGI PhycoCosm [g] | Group 1 (Zn-CDF) |
| No_672670 | *Nannochloropsis oceanica* CCMP1779 | Ochrophyta | Protein Id: 672670 | JGI PhycoCosm [g] | Group 1 (Zn-CDF) |
| No_639981 | *Nannochloropsis oceanica* CCMP1779 | Ochrophyta | Protein Id: 639981 | JGI PhycoCosm [g] | Group 1 (Zn-CDF) |
| No_577472 | *Nannochloropsis oceanica* CCMP1779 | Ochrophyta | Protein Id: 577472 | JGI PhycoCosm [g] | Group 2 (Fe/Zn CDF) |
| No_630732 | *Nannochloropsis oceanica* CCMP1779 | Ochrophyta | Protein Id: 630732 | JGI PhycoCosm [g] | Group 3 (undefined) |
| No_556631 | *Nannochloropsis oceanica* CCMP1779 | Ochrophyta | Protein Id: 556631 | JGI PhycoCosm [g] | Group 4 (Mn-CDF) |
| Osp_404893 | *Ochromonadaceae* sp. CCMP2298 | Ochrophyta | Protein Id: 404893 | JGI PhycoCosm [g] | Group 1 (Zn-CDF) |
| Osp_402386 | *Ochromonadaceae* sp. CCMP2298 | Ochrophyta | Protein Id: 402386 | JGI PhycoCosm [g] | Group 2 (Fe/Zn CDF) |

| | | | | | |
|---|---|---|---|---|---|
| Osp_447574 | *Ochromonadaceae* sp. CCMP2298 | Ochrophyta | Protein Id: 447574 | JGI PhycoCosm [g] | Group 2 (Fe/Zn CDF) |
| Osp_452074 | *Ochromonadaceae* sp. CCMP2298 | Ochrophyta | Protein Id: 452074 | JGI PhycoCosm [g] | Group 5 (undefined) |
| Osp_483158 | *Ochromonadaceae* sp. CCMP2298 | Ochrophyta | Protein Id: 483158 | JGI PhycoCosm [g] | Group 5 (undefined) |
| Osp_198761 | *Ochromonadaceae* sp. CCMP2298 | Ochrophyta | Protein Id: 198761 | JGI PhycoCosm [g] | Unassigned |
| Osp_422041 | *Ochromonadaceae* sp. CCMP2298 | Ochrophyta | Protein Id: 422041 | JGI PhycoCosm [g] | Unassigned |
| Bn_83139 | *Bigelowiella natans* CCMP2755 | Chlorarachniophyta | Protein Id: 83139 | JGI PhycoCosm [g] | Group 2 (Fe/Zn CDF) |
| Bn_89846 | *Bigelowiella natans* CCMP2755 | Chlorarachniophyta | Protein Id: 89846 | JGI PhycoCosm [g] | Group 2 (Fe/Zn CDF) |
| Bn_74255 | *Bigelowiella natans* CCMP2755 | Chlorarachniophyta | Protein Id: 74255 | JGI PhycoCosm [g] | Group 3 (undefined) |
| Bn_34339 | *Bigelowiella natans* CCMP2755 | Chlorarachniophyta | Protein Id: 34339 | JGI PhycoCosm [g] | Group 4 (Mn-CDF) |
| Bn_76194 | *Bigelowiella natans* CCMP2755 | Chlorarachniophyta | Protein Id: 76194 | JGI PhycoCosm [g] | Group 5 (undefined) |
| Bn_88808 | *Bigelowiella natans* CCMP2755 | Chlorarachniophyta | Protein Id: 88808 | JGI PhycoCosm [g] | Group 5 (undefined) |
| Bn_135221 | *Bigelowiella natans* CCMP2755 | Chlorarachniophyta | Protein Id: 135221 | JGI PhycoCosm [g] | Group 5 (undefined) |

[a] www.ncbi.nlm.nih.gov/protein/
[b] genome.jgi.doe.gov/portal/
[c] www.yeastgenome.org
[d] www.arabidopsis.org
[e] mycocosm.jgi.doe.gov/mycocosm/home
[f] www.plantmorphogenesis.bio.titech.ac.jp/~algae_genome_project/klebsormidium/
[g] phycocosm.jgi.doe.gov/phycocosm/home
[h] phytozome.jgi.doe.gov/pz/portal.html

**Supplementary Table S2**. Oligonucleotide PCR primer sequences used for full-length amplification of *C. reinhardtii* MTP cDNAs. Underlined sequence indicates restriction enzyme site used for cloning.

| Gene | Primer name | Sequence (5′ – 3′) |
|---|---|---|
| *MTP1* | MTP1XbaIF | AAATCTAGAAAAATGTCAGAAAGAACGCCTCTTC |
| | MTP1SacIR | AAAGAGCTCCTAGACCTGTGCGTCGCCAGCGGC |
| *MTP2* | MTP2BamHIF | AAAGGATCCAAAAATGGCGCAATTAGCTCGTGAAG |
| | MTP2XbaIR | AAATCTAGAAACTGGGCTGCAAACGGGCTACTTC |
| *MTP3* | MTP3BamHIF | AAAGGATCCAAAAATGGCTCAGTTAGCACGCGAAG |
| | MTP3XbaIR | AAATCTAGACAGCCTACTGTGCCGCGGCGCCAG |

**Supplementary Table S3**. Oligonucleotide PCR primer sequences used for qPCR for *C. reinhardtii* MTP genes.

| Gene | Primer name | Sequence (5′ – 3′) |
|---|---|---|
| *MTP1* | MTP1F | CGTGTGGCTTGAGCGAAGAG |
| | MTP1R | GCTTGCGTGCGATGATACGA |
| *MTP2* | MTP2F | ATGAGTGTGCGGGAGTCGCA |
| | MTP2R | GGCAGTGGCTTCATCACGTC |
| *MTP3* | MTP3F | ATCGAGGCTCTTGACACTGT |
| | MTP3R | GTAAGGCGCTGCTCAGCGTC |
| *MTP4* | MTP4F | ACATGTGTGTGCGGGAGTCG |
| | MTP4R | CTTGTGCCGGTGCAGGGACC |
| *MTP5* | MTP5F | CCTTCATCCTCAGCCATAGCA |
| | MTP5R | CGAGAGGTTGGGAAGTGAG |
| *CBLP* | CBLPF | CTTCTCGCCCATGACCAC |
| | CBLPR | CCCACCAGGTTGTTCTTCAG |

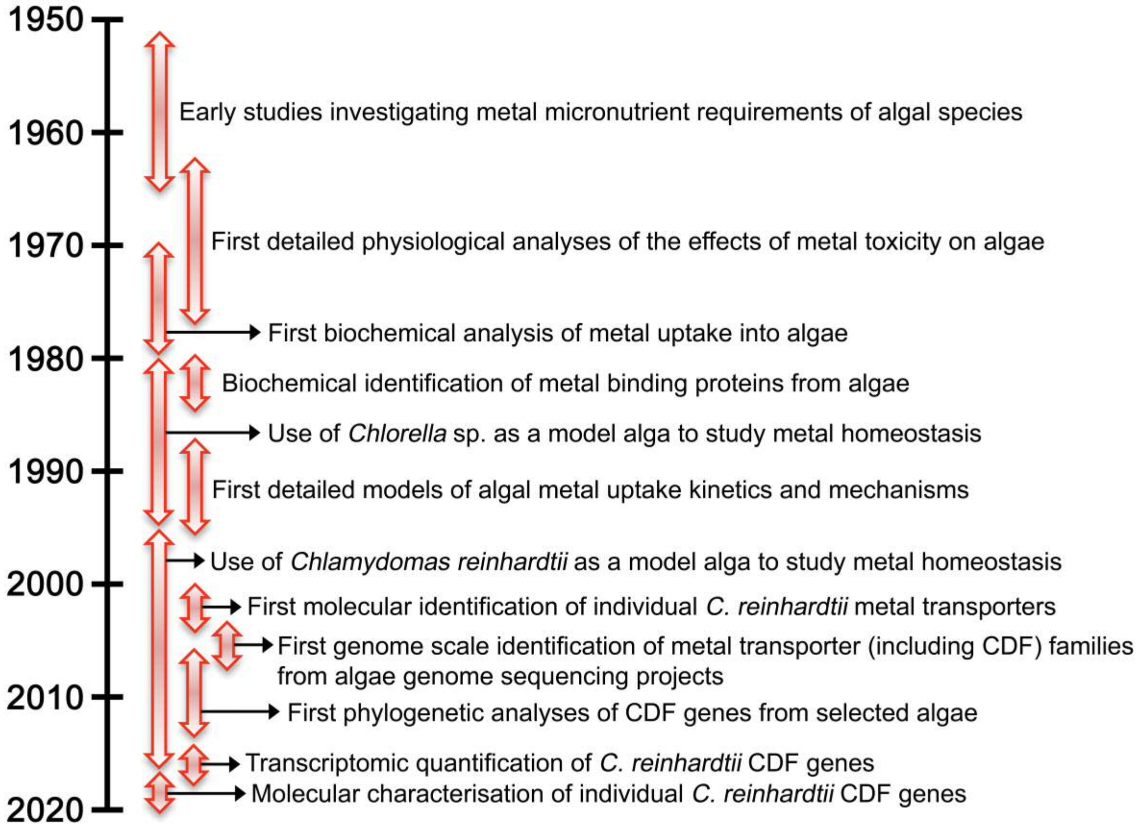


**Supplementary Figure S1.** A timeline of key events in algal metallomics and CDF transporter research over the last 70 years.

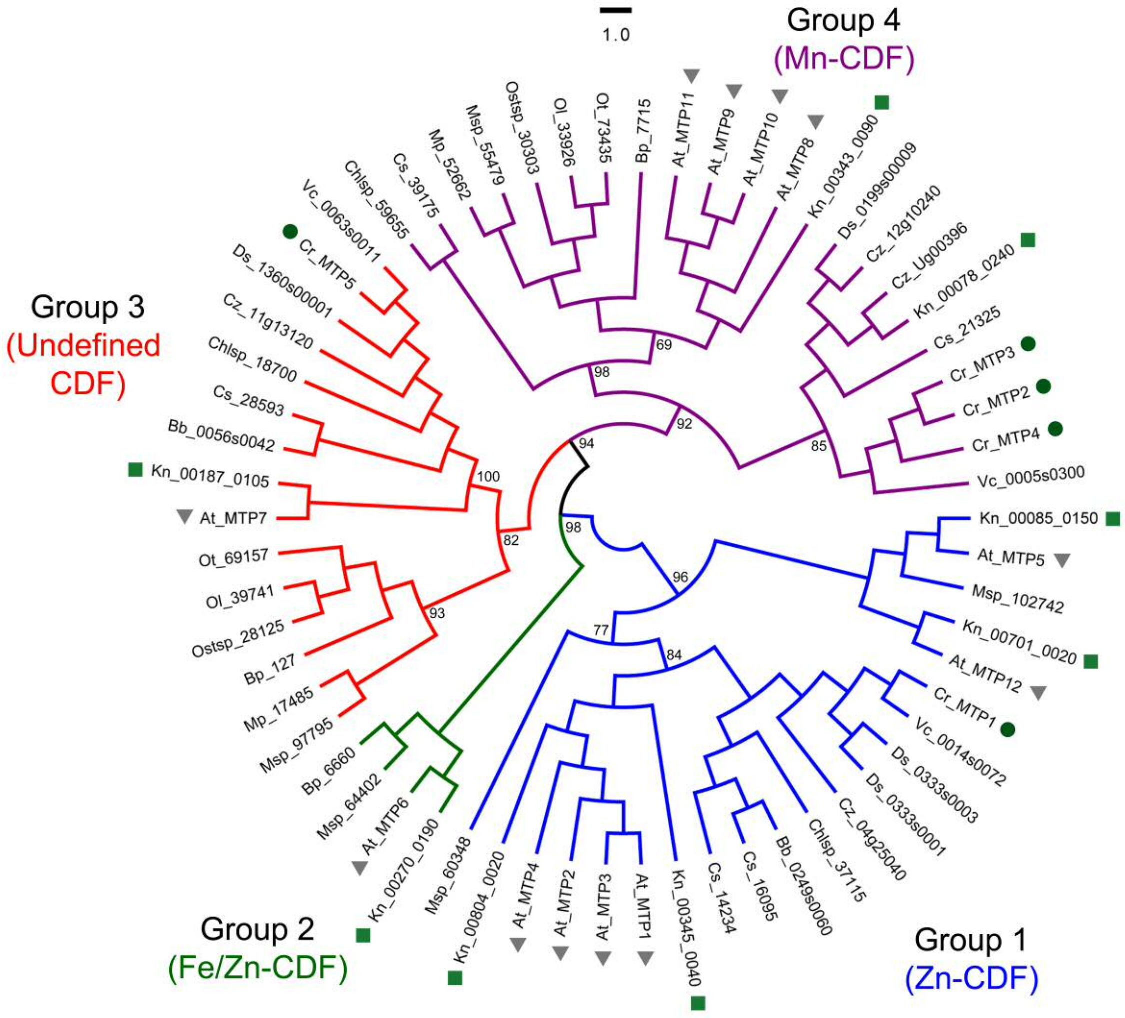


**Supplementary Figure S2.** Phylogenetic analysis of CDF transporter family proteins from chlorophyta algae. CDF protein sequences from the charophyte *Klebsormidium nitens* (square symbols) and land plant *Arabidopsis thaliana* (triangle symbols) were included for comparison. The *Chlamydomonas reinhardtii* sequences are highlighted (circle symbols). The genome identifier or accession numbers of all sequences used are provided in Table S1. Four major groups are highlighted. The tree was constructed using the maximum likelihood method and derived from alignments of full length amino acid sequence. A consensus tree following 1000 bootstrap replications is shown, with bootstrap percentage values indicated at the nodes of major branches. The branch length scale bar indicates the evolutionary distance of one amino acid substitution per site.

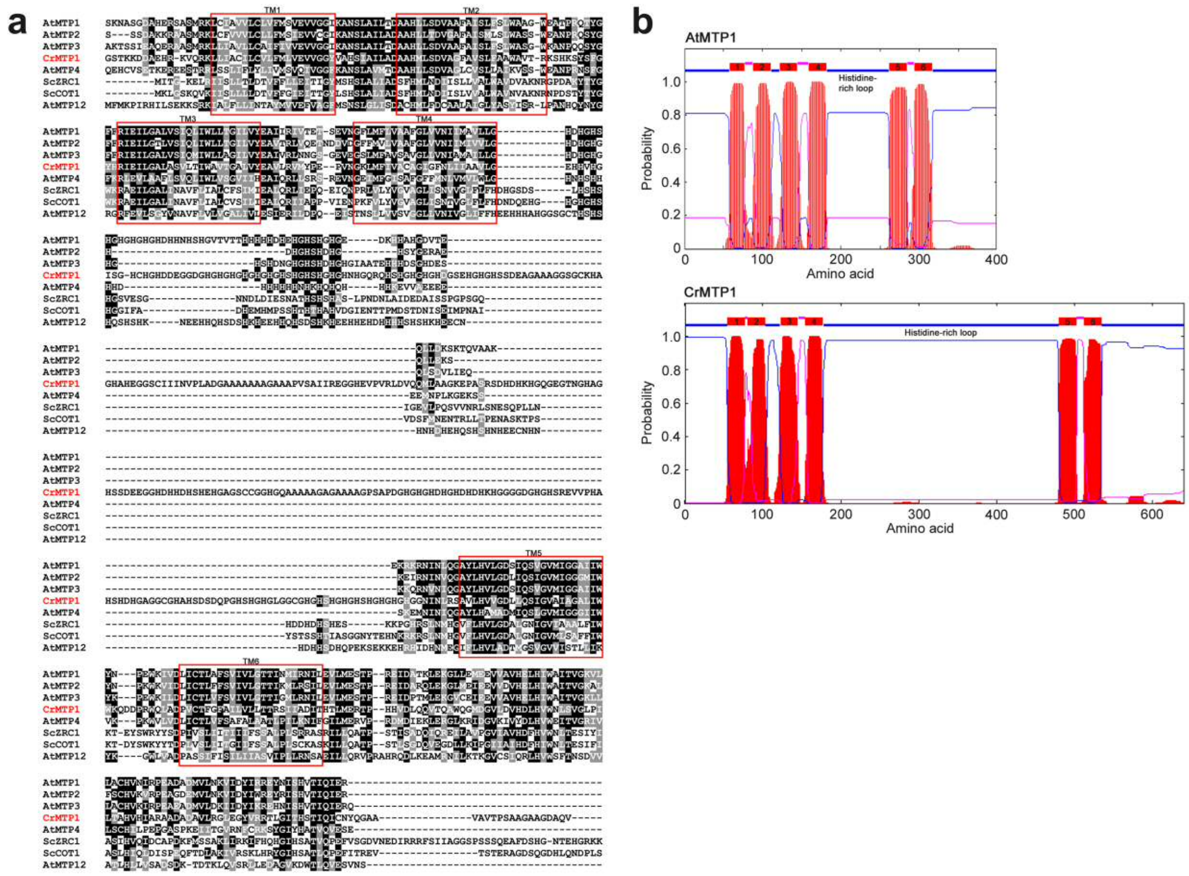


**Supplementary Figure S3.** Sequence analysis of *Chlamydomonas reinhardtii* MTP1 with other Group 1 Zn-CDF proteins. (a) Partial-length multiple sequence alignment of CrMTP1 in comparison with AtMTP1, AtMTP2, AtMTP3, AtMTP4, AtMTP12, ScZRC1 and ScCOT1. The N-terminal tail sequences of CrMTP1 and the *Arabidopsis* MTPs, and the N- and C-terminal tail sequences of the yeast ZRC1/COT1 are not shown. The sequence of CrMTP1 from amino acids 39 and AtMTP1 from amino acid 41 are shown. Amino acids that are identical or similar are shaded in black or grey, respectively. The six putative transmembrane (TM) spans are shown as red boxes and determined by hydropathy analysis (b). (b) Topology models of CrMTP1 in comparison with AtMTP1, generated by TMHMM. Red areas indicate the predicted TM spans.
